# Dynamics of data availability in disease modeling: An example evaluating the trade-offs of ultra-fine-scale factors applied to human West Nile virus disease models in the Chicago area, USA

**DOI:** 10.1101/2021.03.29.437618

**Authors:** J.A. Uelmen, P. Irwin, W.M. Brown, S. Karki, M.O. Ruiz, B. Li, R.L. Smith

## Abstract

**Background:** Since 1999, West Nile virus (WNV) has moved rapidly across the United States, resulting in tens of thousands of human cases. Both the number of human cases and the minimum infection rate (MIR) in vector mosquitoes vary across time and space and are driven by numerous abiotic and biotic forces, ranging from differences in microclimates to socio-demographic factors. Because the interactions among these multiple factors affect the locally variable risk of WNV illness, it has been especially difficult to model human disease risk across varying spatial and temporal scales. Cook and DuPage Counties, comprising the city of Chicago and surrounding suburbs, experience some of the highest numbers of human neuroinvasive cases of WNV in the United States. Despite active mosquito control efforts, there is consistent annual WNV presence, resulting in more than 285 confirmed WNV human cases and 20 deaths from the years 2014-2018 in Cook County alone.

**Methods:** A previous Chicago-area WNV model identified the fifty-five most high and low risk locations in the Northwest Mosquito Abatement District (NWMAD), an enclave ¼ the size of the combined Cook and DuPage county area. In these locations, human WNV risk was stratified by model performance, as indicated by differences in studentized residuals. Within these areas, an additional two-years of field collections and data processing was added to a 12-year WNV dataset that includes human cases, MIR, vector abundance, and land-use, historical climate, and socio-economic and demographic variables, and was assessed by an ultra-fine-scale (1 km spatial × 1 week temporal resolution) multivariate logistic regression model.

**Results:** Multivariate statistical methods applied to the ultra-fine-scale model identified fewer explanatory variables while improving upon the fit of the previous model. Beyond MIR and climatic factors, efforts to acquire additional covariates only slightly improved model predictive performance.

**Conclusions:** These results suggest human WNV illness in the Chicago area may be associated with fewer, but increasingly critical, key variables at finer scales. Given limited resources, these findings suggest large variations in model performance occur, depending on covariate availability, and provide guidance in variable selection for optimal WNV human illness modeling.

## Introduction

West Nile virus (WNV; Family *Flaviviridae*), a mosquito-borne disease originating from the West Nile region of Uganda, first arrived to the United States (U.S., New York, NY) in 1999. Once arriving in New York, the virus took only three years to traverse the contiguous U.S., reaching California in 2002 (1). The virus has now become one of the most widespread arboviruses in the world, and is present in every continent except Antarctica (2). In the Midwestern U.S., mosquitoes of the *Culex* (*Cx.*) genus are the main vectors for transmitting WNV (3). *Culex* mosquitoes are capable of feeding on several hosts to satisfy one blood meal, increasing the opportunity for multiple infections across species (4). Although primarily ornithophilic, prior studies indicate that *Cx.* species may shift feeding preferences to humans later in the summer months (5,6).

From 1999-2018, there have been a total of 50,830 human cases resulting in 2,330 deaths across the US (7). At local scales, drivers of human disease, including WNV, vary in actual effect and magnitude from values reported in studies that more commonly assess disease dynamics at state, regional, or national scales (8). Previous studies have identified similar abiotic and biotic factors associated with human WNV illness, including prior weather conditions (weekly temperature and precipitation lags), mosquito infection and abundance, socio-demographic characteristics of the local population, and level of public awareness and education, but these were all at state or regional scales (9–17).

Karki et al. (2020) and Ruiz et al. (2010) are two of the few studies to evaluate weekly spatiotemporal factors and their associations with human WNV illness at a smaller scale (1-km hexagonal spatial units), in a highly urban 2-county area (Cook & DuPage counties, encompassing the greater Chicago, IL area). This region consistently experiences one of the highest annual WNV incidences in the country (20). While an excellent overall model fit was achieved by using a large number of explanatory variables (n=40), the relative importance of covariates and the resulting strength of disease prediction across the study area varied widely. Understanding how and why these relationships change at specific spatiotemporal locales has been a major conceptual challenge when modeling human WNV illness, and is the central focus for this study.

The Northwest Mosquito Abatement District (NWMAD), occupying the northwest corner of Cook County, is one of Chicago’s four abatement districts responsible for mosquito control, and has an excellent long-term mosquito abundance and testing data throughout its jurisdiction. Human and environmental factors are heterogenous throughout the NWMAD, presenting a strong gradient of human population density, household size and age, socio-economic values, and land-use and land-cover, providing a highly representative enclave of the greater Chicago region.

Specifically, the main objectives of this study were to: (i) evaluate and contrast key variables in this study to the larger Cook and DuPage model, (ii) assess the similarities and differences among locations that were predicted accurately by the larger model and those that were predicted poorly, and (iii) quantify the impact of newly acquired data on prediction of human WNV illness. The authors hypothesize that evaluating human WNV risk at an ultra-fine-scale (UFS) will improve overall model performance (as compared to broader scale models). Additionally, the authors hypothesize that by including several additional covariates that are specific to the UFS study area, the eco-epidemiologic relationships of human WNV transmission will be improved.

## Methods

### Ethics Statement

All data collected from the Illinois Department of Health (IDPH) were through a user agreement approved by the University of Illinois Institutional Review Board and the Illinois Department of Public Health Institutional Review Board. The human activity observation protocol was approved by the University of Illinois Institutional Review Board. Field collections and any use of generated data were approved by the University of Illinois Biosafety Committee.

### Study area

This study was conducted within the NWMAD, a 605-km^2^ area that comprises the northwest suburbs of Chicago (Cook County, IL, Fig 1). The NWMAD study area is an enclave of the Cook & DuPage counties model, the previous research site conducted by Karki et al. (2020). Within the NWMAD study area, fifty-five 1-km hexagonal units were specially selected. These fifty-five 1-km units denote the “ultra-fine-scale” (UFS) study area and contained a total of forty human WNV cases from 2005-2016 (Table S1). By focusing on the spatiotemporal dynamics of WNV transmission in humans in this UFS study area, research efforts have focused on additional data collection, more than doubling the total amount of covariates related to WNV in the Chicago region than the previous Karki et al. (2020) study. Through these additional collection efforts, this study aims to better control, assess, and ultimately, understand the relationships among key predictors of human WNV disease at very fine scales. All model data were summarized and processed within 1-km diameter hexagons, as a neutral configuration in both size and shape, free of any political boundaries. Using statistical selection processes (described below), fifty-five of the 1,019 hexagons within the NWMAD were selected as the observational units for this study.

**Fig 1.**
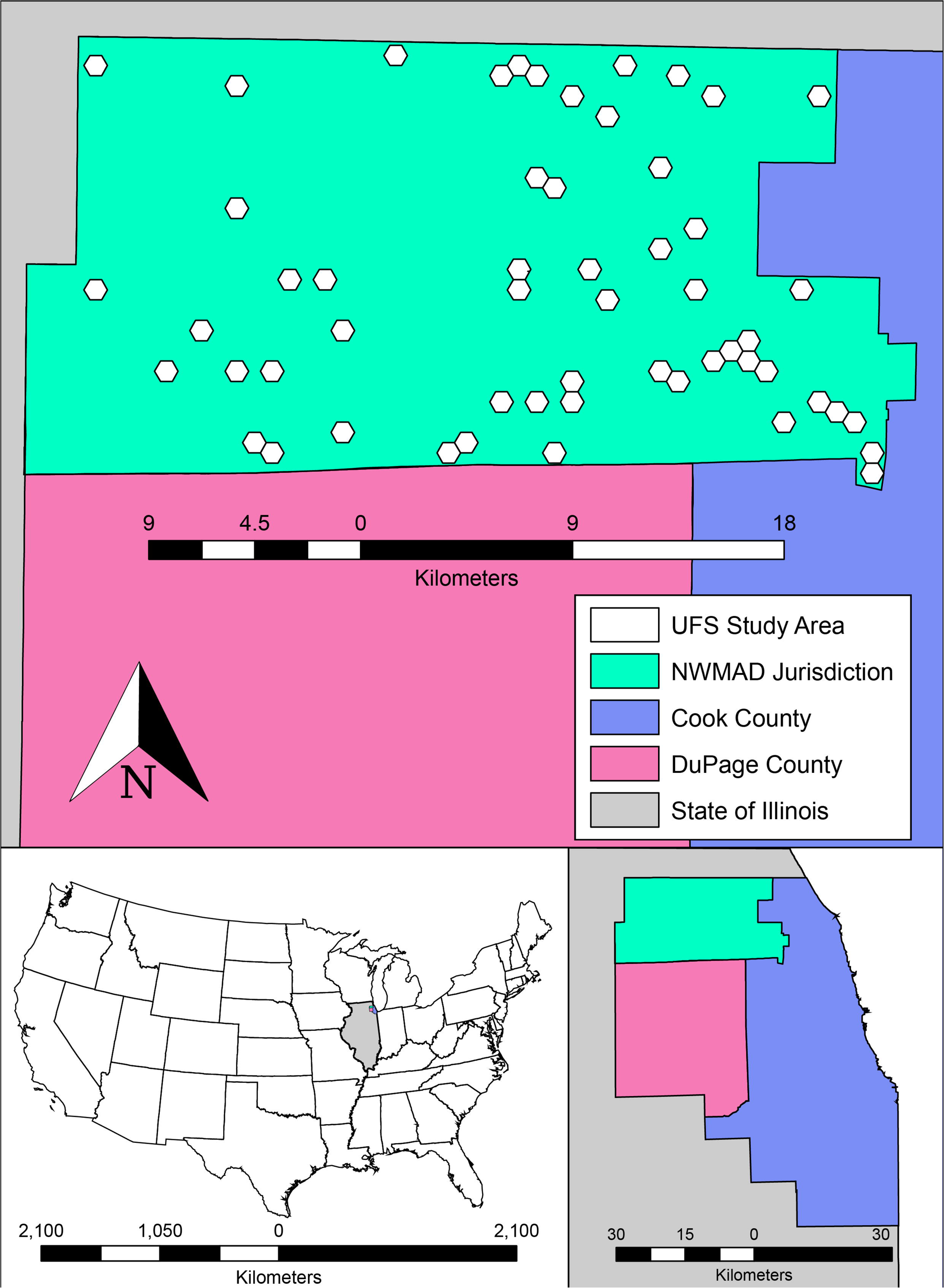
The UFS study area, contained within the Northwest Mosquito Abatement District (NWMAD), in relation to Cook and DuPage Counties. Overlaid are 1-km diameter hexagons, the observational units used in this study. Northwest Mosquito Abatement District comprises 1,019 of the total 5,345 hexagons in all of Cook and DuPage Counties.

### Model covariates

The Cook and DuPage model evaluated forty covariates derived from a variety of abiotic and biotic factors associated with human WNV transmission, including climate and weather records, mosquito infection, environmental land use, and socio-demographic census data. For this study, additional data processing and field collections resulted in forty-two additional independent variables, each determined to be ecologically- or epidemiologically-related to human WNV illness in our study areas of focus (Table 1). Each variable was independently calculated by hexagon and averaged for each Centers for Disease Control and Prevention (CDC) epidemiological week (18-38, Sunday-Saturday) of the years 2005 through 2016 (7). Previously collected data used in this study are explained in detail in Karki et al (2020) and can also be found in supplemental materials.

**Table 1.**
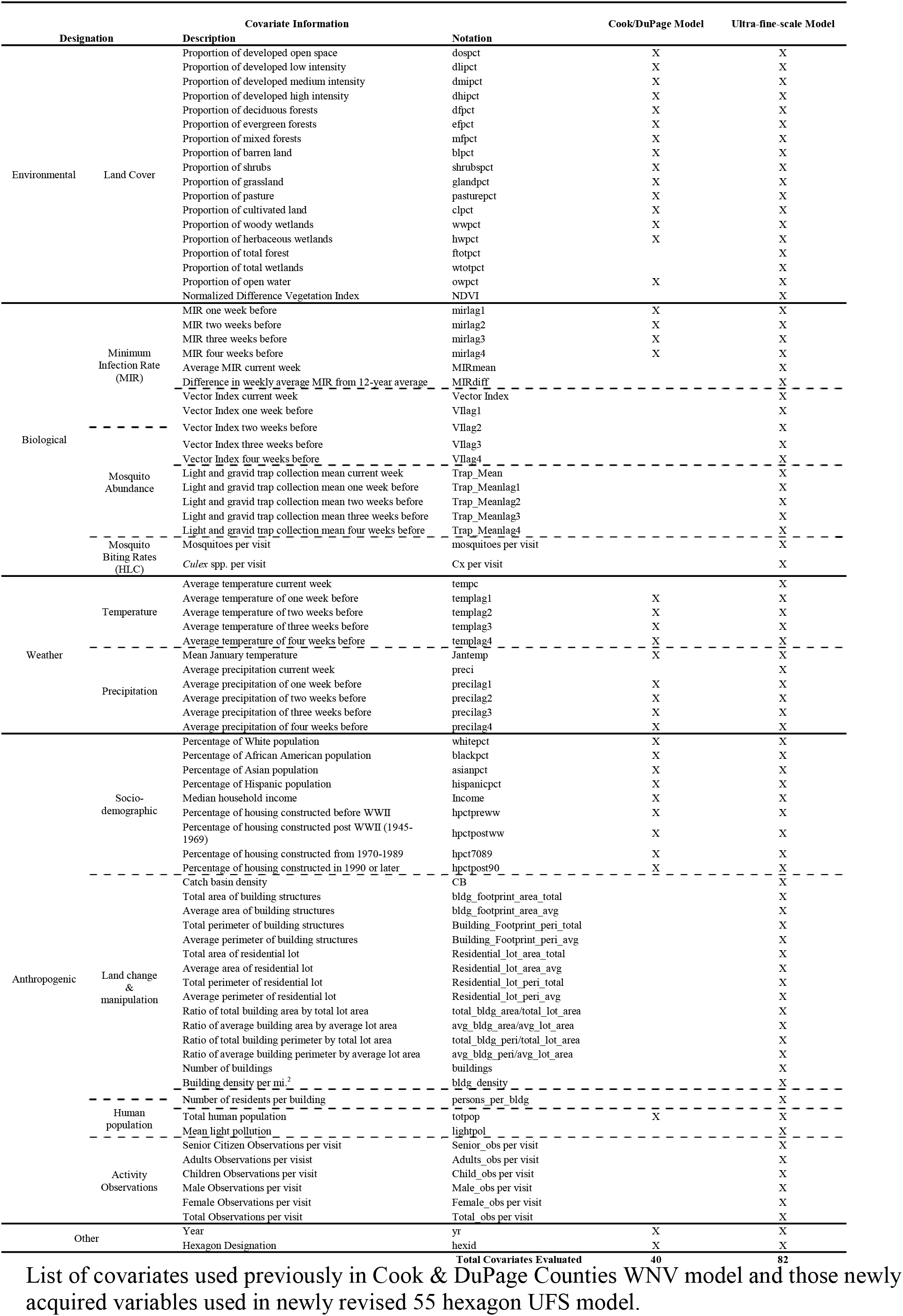
Ecologically- and epidemiologically-related human WNV illness variables assessed.

### Previously existing data

#### Human illness

Human WNV cases in Illinois were classified as either confirmed or probable, as reported to the IDPH by public health or licensed medical professionals (mandatory reporting of WNV cases is required in the state). Human cases were converted into binary form (presence/absence of illness) and weekly case rate, controlling for human population, for each hexagon.

#### Abiotic Predictors

Thirty meter resolution land cover from the 2011 United States Geological Survey (21) National Land Cover Database (NLCD) provided 30 m resolution classified raster data for the NWMAD. There were 15 unique land cover types, ranging from various forests and vegetation to built up urban space. Weekly mean temperatures and weekly precipitation totals, acquired from the PRISM Climate Group (22), were extracted for each hexagon in this study using ArcGIS 10.5.1(23).

### Newly added data

#### Abiotic Predictors

##### Catch basin density

Due to the high preference for breeding in catch basins (e.g. sewers) by *Culex pipiens*, the density of catch basins 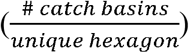 was calculated and assessed. The NWMAD provided point data for each catch basin within its jurisdiction. All point data were then aggregated to each hexagon using the spatial location join feature in ArcGIS. A combined total of 8,443 catch basins were recorded among all hexagons (min = 1, max = 543).

##### Building and residential structures

Previous WNV studies in the Chicago area found a link between the density, size, and age of housing and human cases (9). Through high-resolution (1m) aerial imagery from ArcGIS and USDA (2018), every permanent structure (e.g. residence, shed, garage, deck) was traced and converted to polygons in ArcGIS. The area and perimeter of each polygon was calculated and aggregated for each hexagon. Commercial and residential lots were provided by Cook County Data Catalog (2019), using 2016 tax appropriations. In total, there were a combined 22,892 lots with 24,468 buildings or permanent structures.

##### Light pollution

A recent study by Kernbach et al. (2019) has linked increases in light pollution to WNV in the environment. Because the NWMAD consists of a metropolitan area with an abundance of artificial light, pollution values were evaluated. Light pollution was provided by the New World Atlas of Artificial Night Sky Brightness (27,28). Light pollution was acquired from 2014 data of the VIIRS DNB sensor on the Suomi National Polar-orbiting Partnership satellite. Pixel resolution was 0.75 km; mean value for each 1-km hexagon was calculated in ArcGIS.

#### Biotic Predictors

##### Historical mosquito abundance

The NWMAD consistently collected and diligently maintained their mosquito trapping and identification data throughout the study period. Once deployed, traps were usually checked at least twice a week. Over the 2005-2016 study period, there were a total of 59 traps used in the NWMAD, resulting in a total of 48,406 female *Culex.* spp. from 22 light traps, and 1,110,024 from 37 gravid traps. Weekly mosquito collections by trap were geocoded and interpolated across all hexagons via IDW and extracted using the zonal statistics as table function for each hexagon in ArcGIS. The regular maintenance, collection, and identification, frequency of mosquitoes caught, and distribution of traps within the NWMAD provided strong evidence that mosquitoes collected were representative for the remainder of the study area. Additionally, standard error values as a result of IDW methods were very low, and thus, the assumption for spatial dependency is satisfied (S1 Fig). Mosquito abundance was calculated as the weekly cumulative number of captured female *Culex* spp. from each respective gravid trap (GT) and light trap (LT). Since *Cx. pipiens* and *Cx. restuans* are very difficult to morphologically identify, and with prior studies establishing these as the major *Culex*. species present in this area, all collected specimens from the genus *Culex* were pooled.

##### Normalized Difference Vegetation Index (NDVI)

Trees and shrubs are a major source of nectar and serve as resting places for mosquitoes, especially those that recently blood-fed (29). To evaluate the magnitude of all vegetation, NDVI was incorporated by hexagon, recorded as an average value at three timepoints of each year: CDC epidemiologic weeks 21 (3^rd^-4^th^ week of May), 28 (2^nd^-3^rd^ week of July), and 35 (4^th^ week of August-1^st^ week of September). These CDC epidemiologic weeks mark the center of each the three 8-week active WNV periods in the Midwest, represented as T1 = low WNV activity, T2 = high WNV activity, and T3 = moderate WNV activity. The best available Landsat 7 or 8 bands for each respective time period were acquired from EarthExplorer (30) and processed in ArcGIS.

##### Human activity observations

To provide the most complete measurement of human risk to potential mosquito vectors in nature, this study attempted to quantify human exposure during crepuscular time periods. Human activity observations were conducted in public spaces inside each hexagon, during the crepuscular hours between 6-9:30pm, the preferred feeding period for *Cx. pipiens/restuans*. Observations were conducted within each hexagon for a total of ten minutes per visit. Specifically, a researcher remained stationary for 2 minutes, walked 2 minutes, remained stationary in the new position for 2 minutes, walked back to origination point for 2 minutes, then remained stationary in the original position for 2 final minutes. Human exposure was determined as any period in time a person was outside of any building, vehicle, or enclosed dwelling during the observation period. Observations were classified by apparent gender and age (child, adult, or senior citizen).

##### Human landing catch (HLC)

In conjunction with human activity observations, the number of human-seeking mosquitoes that attempted to blood-feed were collected via human landing catch methods for a fifteen-minute period at each hexagon weekly. To mitigate actual biting events, the researcher would expose only one limb (arm or leg) at a given time. Any mosquito that landed was collected via mechanical aspirator and transferred to a 2 ml collection vial. All collected mosquitoes were transported to the NWMAD within 2 hours and stored at −80°C. All mosquito specimens were identified to species within three days. Any mosquitoes identified as *Culex* spp. were sent to the Fritz Lab at the University of Maryland for species confirmation by *Cx. pipiens* group-specific primers via PCR.

##### Vector Index

The vector index (VI) was calculated as an estimate of the relative number of WNV-infected mosquitoes. For this study, VI was calculated as the average number of pooled *Culex* spp. collected per trap-week multiplied by the proportion of mosquitoes infected with WNV. The following equation was modified from the CDC (2013):

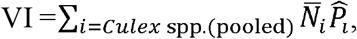

where 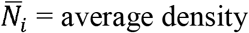 (number of mosquitoes per trap week) and 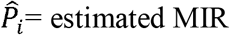 (proportion of mosquito pools testing positive for WNV). Calculated weekly VI for each trap by week was then interpolated via IDW method for estimations across the NWMAD.

##### Nuisance Factor and Human WNV Added Risk

The combination of human activity observations, serving as a proxy for potential mosquito bloodmeals, and HLC data, serving as a proxy for potential rate of mosquito biting, formed two unique WNV disease indices: the Nuisance Factor and Human WNV Added Risk. Since the majority of mosquitoes collected were non-*Culex*, a quantitative index, nuisance factor, was created to provide a risk spectrum of encountering nuisance mosquitoes in a given hexagon. The following equation defines the nuisance factor:

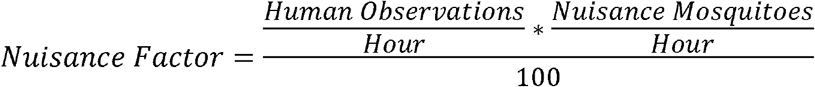

Nuisance factor values ranged from a low of 0 to a high of 32.3. To quantitatively estimate potential risk for exposure to disease within a given hexagon, the human WNV added risk factor was created. This index is defined by the following equation:

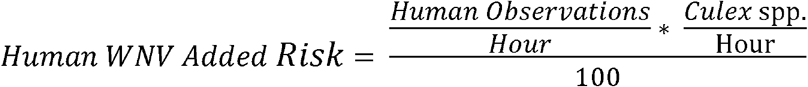

Human WNV added risk ranged from a low of 0 to a high of 1.44.

### Statistical methods

#### Location selection

Of the total 1019 hexagons within the NWMAD, fifty-five (5.4%) were selected as the maximum number of sites that our research team could visit for fifteen minutes each, weekly. The subset of fifty-five hexagons were selected based on two criteria: (1) human population was > 0, and (2) the previous Cook and DuPage model either predicted human WNV extremely well or extremely poorly, as determined by the 2005-2016 average residual output. Furthermore, the residual output was stratified by those locations that had or had not experienced a human case during the 12-year period. These processes created a performance spectrum consisting of five categories of hexagons: negative residuals without a human case (NR0), low residuals without a case (LR0), low residuals with a case (LR1), positive residuals without a case (PR0), and positive residuals with a case (PR1) (Table S1). No hexagons with negative residuals in the Cook and DuPage model had experienced a human case. The spatial arrangements of these hexagons provide adequate coverage of the NWMAD’s jurisdiction (Fig 2).

**Fig 2.**
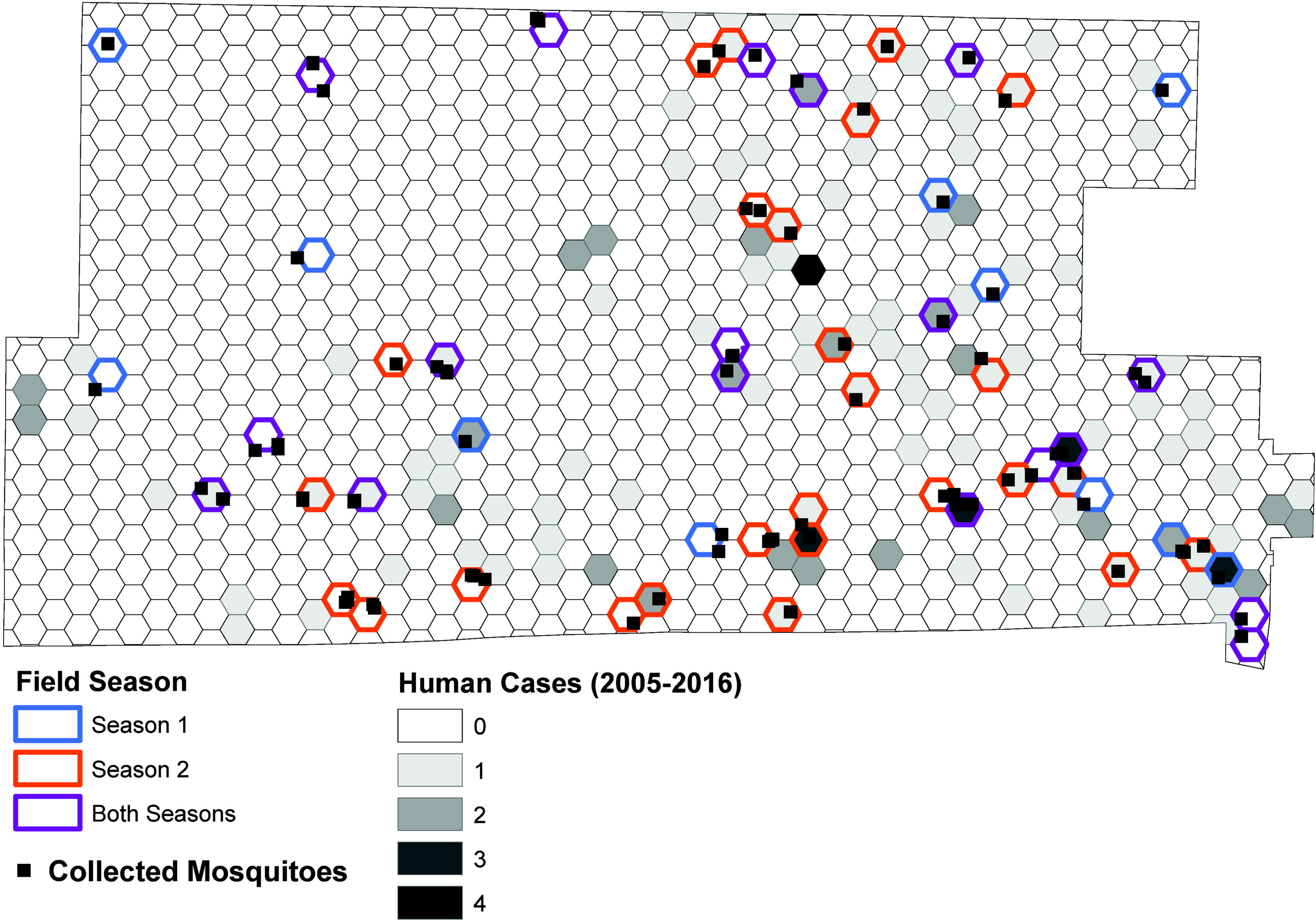
Location of the 55-hexagon study area within the Northwest Abatement District. Hexagons are labeled by field season visited for mosquito collections and human activity observations (color outline) and by total human cases from 2005-2016 (gray scale shaded interior).

#### Model Selection

Two seasons of field collections and processing of new data provided the UFS model with an additional 42 covariates not made available in the previous Cook & DuPage model. The generation of linear and logistic regression models began with a two-step selection process for the initial covariate inclusion: (1) conduct a univariate analysis with each predictor (independent variable) to the WNV disease outcome (binary = logistic, case rate = linear, dependent variable). Candidate variables for multivariate analysis were selected using slightly more conservative p-value than Bursac et al. (2008), p-value ≤ 0.20 vs. ≤ 0.25). Models that create cut-off values of p-value ≤ 0.1 for purposeful univariate covariate selection can erroneously prevent important variables from entering final models (33,34); (2) the final model, a generalized linear model with a Poisson distribution and probit link function, was selected using forward selection method, selecting the final model based on the Bayesian information criterion (BIC). Non-significant covariates were removed from the final model as a product of the iterative selection process. Secondarily, a receiver operating characteristic (ROC) curve was used to visualize overall model performance and Area Under the Curve (AUC) was calculated. All predictors were evaluated for multicollinearity using the PROC REG procedure (SAS Institute Inc. Cary, NC, USA) (S2 Table). Regression analyses were analyzed using the Fit Model feature in JMP 14.2.0 (SAS Institute Inc. Cary, NC, USA). Binary WNV case outcome was analyzed with as a nominal logistic personality. The continuous WNV case rate outcome was analyzed as a standard least squares personality.

#### Model Comparisons

Human WNV illness in the NWMAD was assessed under four model environments, each expressing a defined set of specific parameters. The four model environments were:

1. MIR & Mosquito Abundance (contains no VI covariates),
2. Vector Index (contains no MIR or mosquito abundance covariates),
3. Best-Fit (best fit with all covariates in respective assessment), and
4. Global (all covariates made available in respective assessment)

As a comparison, the original Cook & DuPage model (Karki et al. 2020) was fit using only 40 covariates. Each of these four model environments were assessed using four different covariate sets:

1. All covariates (82 available covariates),
2. Excluding HLC and human observations covariates (74 available covariates),
3. Force-fitting HLC and human observations covariates (8 forced covariates, 82 available covariates), and
4. Only the covariates made available to the Cook & DuPage 2019 model (control model, 40 available covariates).

Under each model environment and covariate set, the outcome of human WNV illness was analyzed using:

1. Logistic regression (presence/absence human WNV illness) and
2. Linear regression (WNV case rate) methods.

In total, there were 36 models assessed (Fig 3); models are named using the convention E_x_C_y_O_z_, where x is the model environment number (0-4, with number 0 assigned to the control environment), C is the covariate set number (1-4), and O is the outcome number (1-2). For both logistic and linear regression, each of the four model environments was fit using each of the four covariate sets. In addition, the control models using only the covariates from the final Cook & DuPage model applied to the UFS region were fit with and without force fitting HLC and human observation covariates.

**Fig 3.**
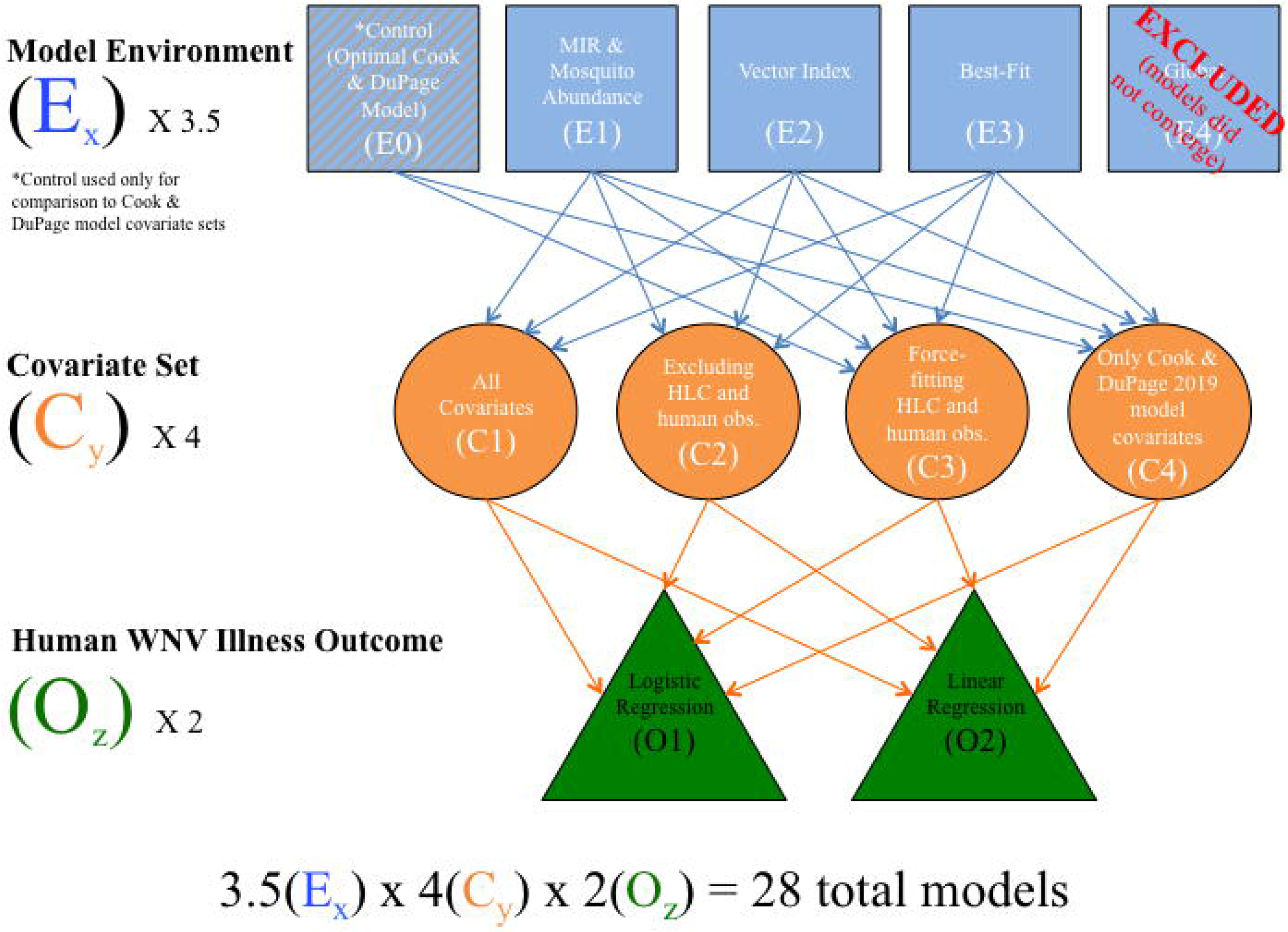
Flow diagram displaying how models were characterized, assembled, and compared in this study. Global models failed to converge and were excluded from the final results. The control model (optimal Cook & DuPage Counties (2019) model) was only used as a comparison for covariates made available only to that original model. Of the original 36 models initially assessed, 8 were removed, resulting in 28 final models assessed in this study.

Half of the models were assessed under logistic and linear outcomes, respectively, and based on the *# of Significant Covariates* (quantity of variables included in final model with p<0.05) and *Degrees of Freedom* (the number of values in the final model that are free to vary). Overall model performance was determined by BIC. While BIC and Aikake’s Information Criterion (AIC) are both maximum likelihood estimators, BIC was chosen to determine model strength due to its stronger penalty term for covariate inclusion (35).

#### Covariate Performance

Similarly to the model performance index, to evaluate the performance for all covariates across 18 logistic and 18 linear models, each of the 82 covariates were standardized by creating the following index:

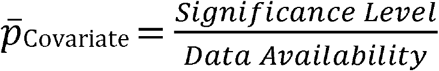

where: *Significance Level* = significance level of covariate in each of the 36 final models (p<0.001 = 4, p<0.01 = 3, p<0.05 = 2, included in the final model = 1), and *Data Availability* = resources required to acquire a respective covariate (level 1 = data widely available, no processing needed, level 2 = data available, requires minimal to moderate processing/analyses, level 3 = data available, requires extensive processing/analyses, level 4 = data not available, needs to be collected, processed, and analyzed, S3 Table). The final net prediction:availability tradeoff used to create the Data Availability variable are categorical and based on the authors’ personal experiences with data used in this study.

## Results

### Model Comparison

The highest performing WNV human risk models were E_3_C_4_O_3_ (Cook & DuPage Best Fit, df = 8, BIC = −227444) and E_2_C_4_O_1_ (Cook & DuPage + VI, df = 14, BIC = 576.2), for linear and logistic regressions, respectively (S4 & S5 Tables).

The top five models that predicted human WNV cases strongest were represented by the control (E_0_, n=2), best-fit (E_3_, n=2) and vector index (E_2_, n=1) environments (Fig 4B, Table 2). These models’ corresponding covariate sets were represented by variables only available to the original Cook & DuPage models (C_4_, n=4), and force-fitting HLC covariates (C_3_, n=1) environments.

**Fig 4.**
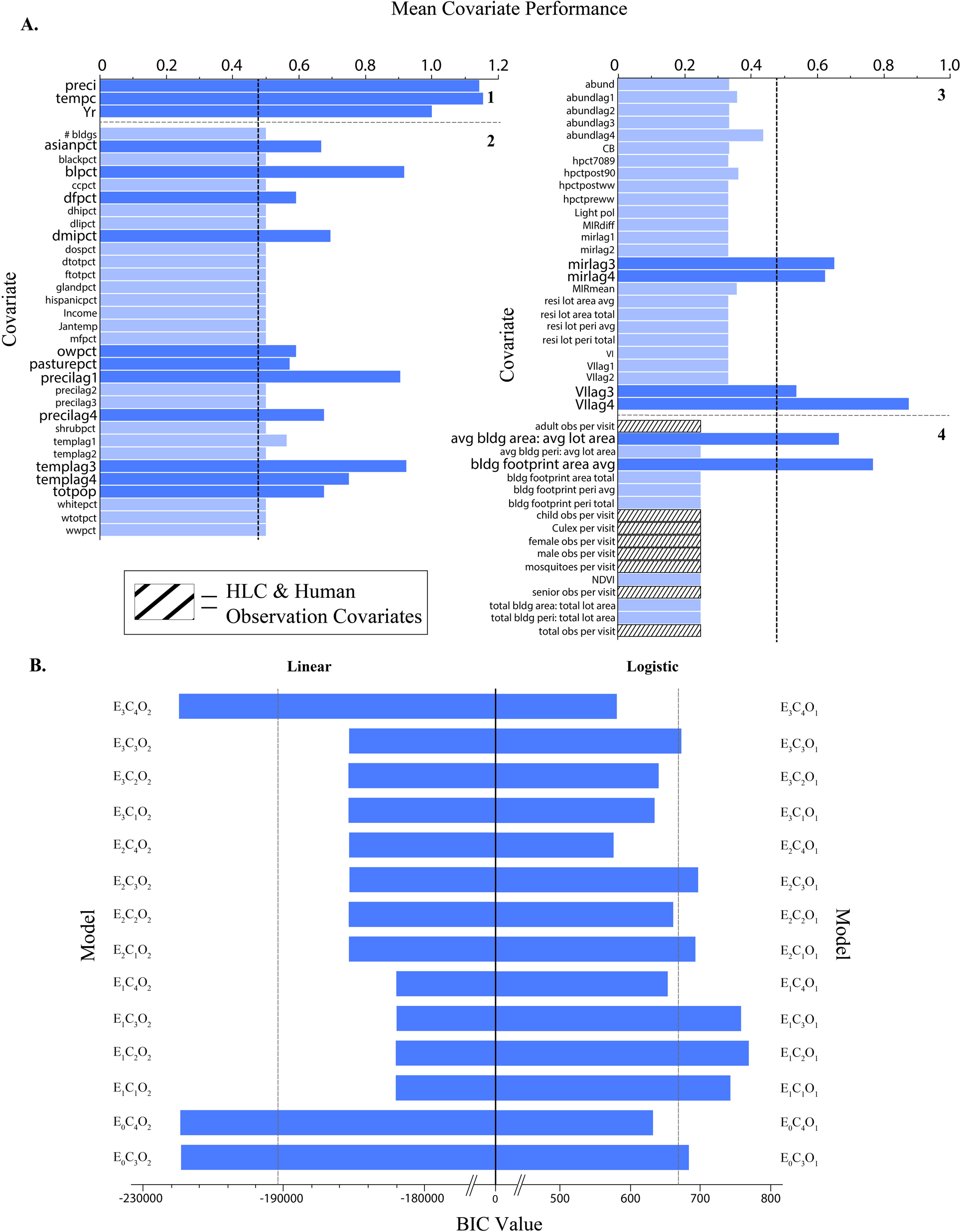
Overall performance for each predictor and final model used in this study. Each of the 70 covariates used in the study, listed in alphabetic order by data availability/work load to acquire score (1-4), were evaluated by mean performance (A). Highest performing covariates are noted by enlarged label text and darker blue bar color. The overall performance for each linear and logistic model (n=14 for both) was evaluated by BIC value (B). Means for each outcome (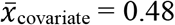; 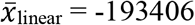; 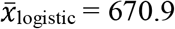) are designated by vertical dashed lines. Details of scoring for each covariate and model are provided in 3 & S3 Tables.

**Table 2.**
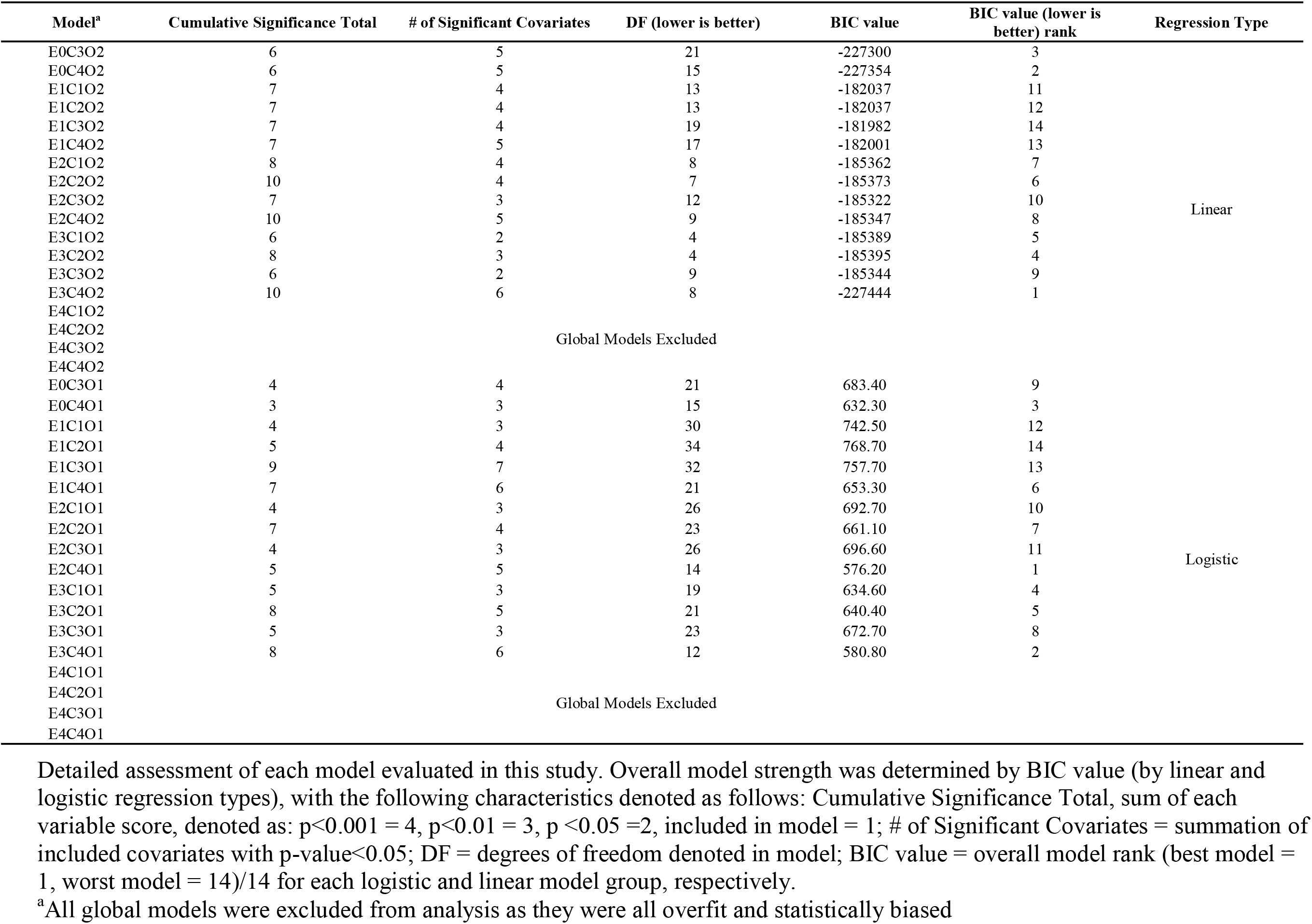
Overall assessment of performance for each model environment, covariate, and outcome combination.

### Covariate Performance

Of the 82 available covariates, 70 (85.4%) were included at least once among a given model, excluding the overfit global models (individual predictor summaries located in S6 Table). Of the 41 covariates (58.6%) that were greater than the mean covariate performance, seven were highly efficient (determined by natural break in the distribution), providing a crude estimation as most valuable variables for human WNV estimation (Fig 4A). These covariates are provided here in descending order of most importance: tempc (temperature (°C), 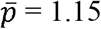), preci (precipitation (mm), 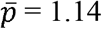), Yr (year, 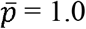), templag3 (temperature lagged by 3 weeks, 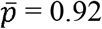), blpct (barren land (%), 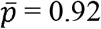), precilag1 (precipitation lagged by 1 week, 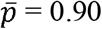), and VIlag4 (vector index 4 weeks prior, 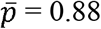). All eight HLC and human observation covariates were included in at least one final model, but none performed highly 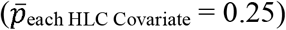. Estimates and calculations for individual covariates are available in S3 Table.

The eight HLC and human observation covariates provided significant differences (*P* ≤ 0.05) in observations and mosquito collections by hexagon type (Figs 5A & 5B). The indices, nuisance mosquito exposure and human WNV added risk, significantly differed by hexagon type (Fig 5C). Hexagons designated as PR1 (positive residual (underpredicted actual cases) with a prior human WNV case) were found to have the most human observations and collected mosquitoes (from both *Culex* and non-*Culex* spp.) per visit. This combination of factors provides hexagons among the PR1 type as the most “risky” in regard to human WNV added risk and increased nuisance mosquito exposure (Fig 5).

**Fig 5.**
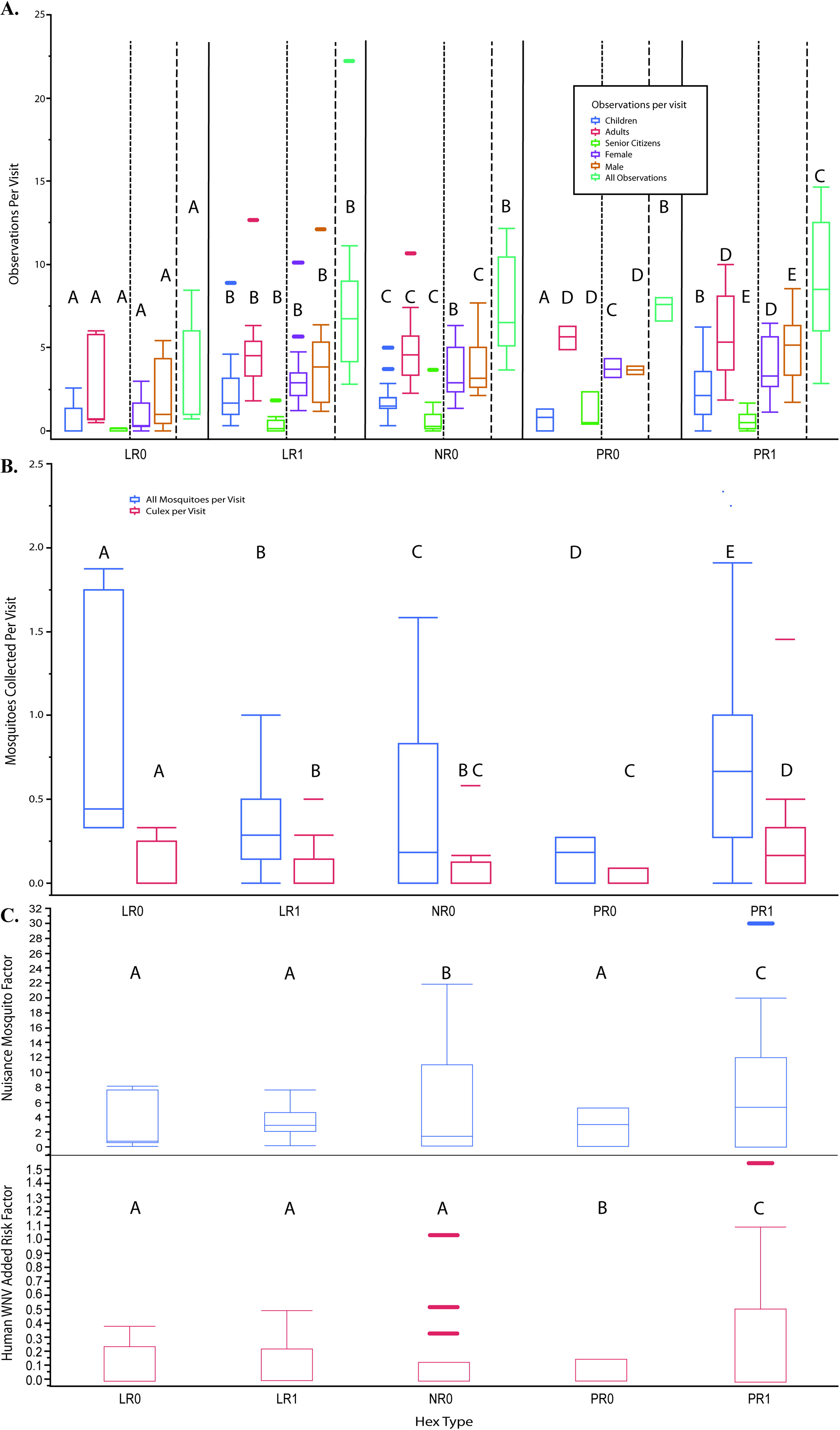
Relationship of hexagon by type. Hexagon type (LR = low residual, PR = positive residual, NR = large, negative residual; 0 = no human case, 1 = human case) are detailed by human observations per visit (A), mosquitoes collected per visit (B), and a product of the two former variables, nuisance factor and WNV added risk (C). Letters above each box and whisker plot designate significantly different groups by hexagon type, as calculated by Tukey’s HSD.

## Discussion

In addition to model comparisons, this study evaluated the performance of the newly acquired VI in comparison to the more commonly used MIR in combination with mosquito abundance. Overall, when fit to the UFS study area, adding mosquito abundance and associated 4-week lags improved this model. When evaluating WNV prediction as a linear outcome, the best-fit model using only covariates available to the original Cook & DuPage model was the highest performing in WNV predictability. However, when evaluating WNV prediction as a binary outcome, VI and its associated 4-week lags replaced MIR as the best predictor of human WNV. While no model emphasizing MIR and abundance was selected as one of the best predictive models, at least one of these variables (and their associated lags) were represented in 4 of the 5 best models (control and best-fit, n=2 for each model). On the contrary, VI, as an emphasized model environment, was selected as the best performing logistic model. Both MIR and VI are critical components in predicting WNV. Under ideal settings, the VI is the preferred method for estimating the risk of mosquito infection, as opposed to MIR. However, deciding between the two biological indicators will be largely dependent upon the data availability for each model of interest. Our study suggests that if resources are limited, net model value leans in favor of using MIR.

The addition of 42 new covariates required a significant allocation of resources but provided minimal benefits towards reducing variance in human WNV prediction. Fortunately, this study suggests that excellent disease prediction models can be achieved with conventional covariates that are publicly available, requiring little to no processing and/or analyses (data availability scores ≤ 2, Fig 4B). However, any covariate used should be adjusted and properly designed for the highest spatial and/or temporal resolution possible, which may require additional efforts to accomplish.

Extensive review of literature indicated no other studies have evaluated covariate strength given limited resources, particularly in the context of making decisions to acquire data. Therefore, the categorizations of covariates by resource allocation (values ranging from 1 (low) to 4 (high)) are based on the experiences of the authors during this study. These values are subjective and may vary across institution or research group, but they may be used as a general estimation in model selection and decision-making. For example, variables related to building and lot size (avg bldg. area: avg lot area, bldg. footprint area avg, bldg. footprint area total, bldg. footprint peri avg., bldg. footprint peri total, and total bldg. area: total lot area) were all ranked a value of 4 because of extensive data processing and review. The authors downloaded high resolution, cloud-free satellite images that were used as a basemap for digital tracing of every building structure (houses, businesses, sheds, detached garages, storage units, etc.) and lots (residential and commercial). This resulted in >47,000 structures and lots digitally traced manually. On the other hand, weather variables (e.g. preci, tempc) were ranked a value of 1 because very little resources were devoted to have the data in a “ready” state. The source of these data, PRISM Climate Group, allows for monthly summaries to be downloaded and extracted with one quick geostatistic process.

This study also aimed to address a key missing index that few studies have evaluated: the relationship of human activity, mosquito exposure, and WNV disease risk. While the related variables did not greatly impact overall model strength, they did provide key insight into a potential key in WNV ecology – the areas that were previously underpredicted with recorded human WNV (hex type: PR1) were consistently found to have the most human activity at crepuscular times, the most mosquitoes overall, and the most *Culex* mosquitoes. However, our results appear to contradict the findings of Read et al. (1994), who discovered that as reports of biting nuisance mosquitoes increased beyond 2 per minute, outdoor human activity rapidly declined. Our results indicate that as mosquito collections increased, human observations also increased (Fig 5). Not only is this a potentially dangerous combination that can foster environments ideal to mosquito-human spillover, previous modeling efforts failed to capture these cases. Future directions will target these highly susceptible locations and aim to capture any additional unaccounted variance.

Like all disease modeling efforts, there are always reporting biases that directly affect true case prevalence. Unfortunately, many vector-borne diseases are largely underreported (37–40), as human cases are vastly overlooked or misdiagnosed, largely due to low severity in disease manifestation in a majority of cases (41,42). This creates difficulties in predicting when and where VBD incidence will arise. Specifically regarding WNV, it is estimated that about 80% of human infections are unreported, as clinical signs are minor or asymptomatic (43,44). The remaining 20% of humans develop West Nile fever, and among this group, about 1% will develop severe and sometimes fatal neuroinvasive disease. In the Chicago area, models in both the UFS and Cook & DuPage locations have very high human WNV prediction capabilities. Despite having among the highest total number of human WNV cases in the U.S. (20), this region has more observational units denoted as non-cases than cases. That has resulted in models with excellent accuracy in predicting where there are no human cases, thus inflating the true accuracy of our models. Nonetheless, while our models are able to reliably predict where human cases are present, the magnitude of effect can be missed (e.g. “hot spots” with greater than 1 case may not be represented).

Disease modelers need to be cognizant of saturating their efforts, both statistically and biologically. Statistically, additional and meaningful covariates will usually improve model fit parameters. However, the inclusion of too many variables can result in overfitting, resulting in models failing to converge (45–47). It is possible that no matter the amount of effort to improve model fit, there is an element of variability attributed with infected humans not seeking medical attention and thus, reducing true disease prevalence (48).

Overall, when compared to the Cook & DuPage model, the best UFS models required fewer predictors and produced a stronger overall fit using most, if not all, the same covariates made available to both model types. Spending the resources (time, money, human-power, processing, analyses, logistic, etc.) to acquire additional covariates may not necessarily be worth the impact on improving human WNV modeling predictions. Rather, fine-tuning the traditional covariates (climatic, weather, and MIR, for example), to the highest spatiotemporal resolution possible may be the most efficient use of resources to minimize variance in VBD prediction models.

## Conclusions

1. The factors and their overall effect on the prediction of human WNV cases differs across scale. Although improved, in comparison to the control Cook & DuPage model applied to the same study region, the “best fit” UFS model AUC = 0.89, suggesting newly unaccounted variances are present.
2. Both vector index and MIR contribute to high performing human WNV prediction models under UFS study areas. In direct comparison, VI is favorable to MIR. However, given limited resources in acquiring and processing additional data, MIR is more efficient for predicting human WNV illness.
3. The effort and resources required to acquire additional covariates, most of which are not publicly available, demonstrate a slight improvement in model prediction and appear less important in reducing variance.
4. In addition to the conventional WNV covariates, namely weather and infection rates, land-use and land-cover and SES/demographic information are widely available with little to no processing or analyses required, and provide the breadth to develop excellent prediction models. However, any covariate utilized must be structured at the finest spatial and/or temporal resolution possible.
5. Human exposure to mosquito biting rates provided minimal benefits to model prediction. More importantly however, these two covariates provided potentially key insight to the susceptibility of humans in locations where WNV is prevalent. Additionally, where WNV is less of a concern, these results provide insight into nuisance mosquito exposure that may lead to improvements in targeted control efforts.

## Acknowledgments

The authors would like to thank Roger Nasci for providing expert insight in vector biology and modeling efforts, Dan Bartlett for providing geospatial data and detailed field information, the Megan Fritz lab for providing confirmatory *Culex* identification, and Chris Stone and Andrew Mackay from the Illinois Medical Entomology lab for providing guidance and expertise with human landing catch methodology. Dr. Marilyn O’Hara Ruiz passed away before the submission of the final version of this manuscript. The corresponding author, Dr. Johnny Uelmen, accepts responsibility for the integrity and validity of the data collected and analyzed.

## Declarations

### Availability of Data and Materials

The dataset supporting the conclusions of this article is available in the University of Illinois repository, https://doi.org/10.13012/B2IDB-5901636_V1.

### Authors’ contributions

JAU conceived the presented idea, collected field samples and provided data analysis and processing. PI provided research assistance and expertise in mosquito collection and biology. SK provided expertise and additional datasets for analysis. WMB provided analytical assistance and provided data sources. BL provided statistical oversight and expertise. MOR provided the initial product idea, planning, and supervision. RLS provided oversight of all aspects throughout the study. All authors discussed the results and contributed to the final manuscript.

### Funding

This publication was supported by Cooperative Agreement #U01 CK000505, funded by the Centers for Disease Control and Prevention. Its contents are solely the responsibility of the authors and do not necessarily represent the official views of the Centers of Disease Control and Prevention or the Department of Health and Human Services.

### Consent for publication

Not applicable

### Competing interests

The authors declare that they have no competing interests.

## Supporting Information

**S1 Fig. Measurement of standard error associated with interpolated** *Culex* **species abundance by light (A) and gravid (B) traps (averaged for all traps), and mean MIR (C) for each of the 55 hexagons, from 2005-2016**. The average weekly mosquito abundance multiplied by average weekly mean MIR created a third infection parameter, the vector index.

**S2 Table. Correlation matrices for each of the 82 covariates assessed in this study.** Tables are grouped by anthropogenic (A), biological (B), environmental (C), weather (D), or other (E), as indicated in Table 1.

**S3 Table. Detail of scoring and categorization of covariates used across all models assessed in this study, organized by data availability/work load to acquire score.** Scoring was denoted as follows: Cumulative Significance, p<0.001 = 4, p<0.01 = 3, p <0.05 =2, included in model = 1; Data Availability/Work Load to Acquire, Data Unavailable/Fieldwork required = 4, Data available, but requires many resources to use = 3, Data available, but requires moderate resources to acquire = 2, Data readily available and requires little to no resources to use = 1; Covariate Value = Quotient of previous two columns.

**S4 Table. Model fit comparisons of the UFS hexagons, applying (A) newly acquired data (excluding HLC and human observations, covariate set 2), or (B) only the covariates made available to the previously published Cook & DuPage model (covariate set 4).** Each model outcome was assessed using logistic (presence/absence WNV human illness case) and generalized linear (WNV case rates, controlling for human population) methods. Asterisks indicate level of statistical significance (* = p ≤ 0.05, ** = p ≤ 0.001, *** = p ≤ 0.0001

^a^Logistic regression outcome = human WNV presence/absence per hexagon, per week; GLM outcome = WNV human case rate (per hexagon, per week).

^b^ROC applies to only logistic regression

^c^As the final selected model in the Original Cook & DuPage paper (2019), this model environment was assessed only for the comparison to the Cook & DuPage models for this study and not applied to the UFS model. The original model covariates, eftpct and ehwpct, have 0 observations among the selected 55 hexagons and were removed.

**S5 Table. Model fit comparisons of the UFS hexagons, using best-fit models with additional human landing catch and human activity observations to incorporate added human risk.**

Human risk covariates were added to the UFS model by (**A**) best-fit integration (covariate set 1) and (**B**) force-fitting (covariate set 3). Asterisks indicate level of statistical significance (* = p ≤ 0.05, ** = p ≤ 0.001, *** = p ≤ 0.0001

^a^Logistic regression outcome = human WNV presence/absence per hexagon, per week; GLM outcome = WNV human case rate (per hexagon, per week).

^b^ROC applies to only logistic regression

^c^As the final selected model in the Original Cook & DuPage paper (2019), this model environment was assessed only for the comparison to the Cook & DuPage models for this study and not applied to the UFS model. The original model covariates, eftpct and ehwpct, have 0 observations among the selected 55 hexagons and were removed.

**S6 Table. Mean and standard error values for each predictor evaluated in this study.**

Values represent averages for all hexagons (pooled) over the entire study period (2005-2016).

## Notes

### Competing Interest Statement

The authors have declared no competing interest.

