## Supplementary figures and images for "Dynamics of data availability in disease modeling: An example evaluating the trade-offs of ultra-fine-scale factors applied to human West Nile virus disease models in the Chicago area, USA"

### Supplemental Figure 1

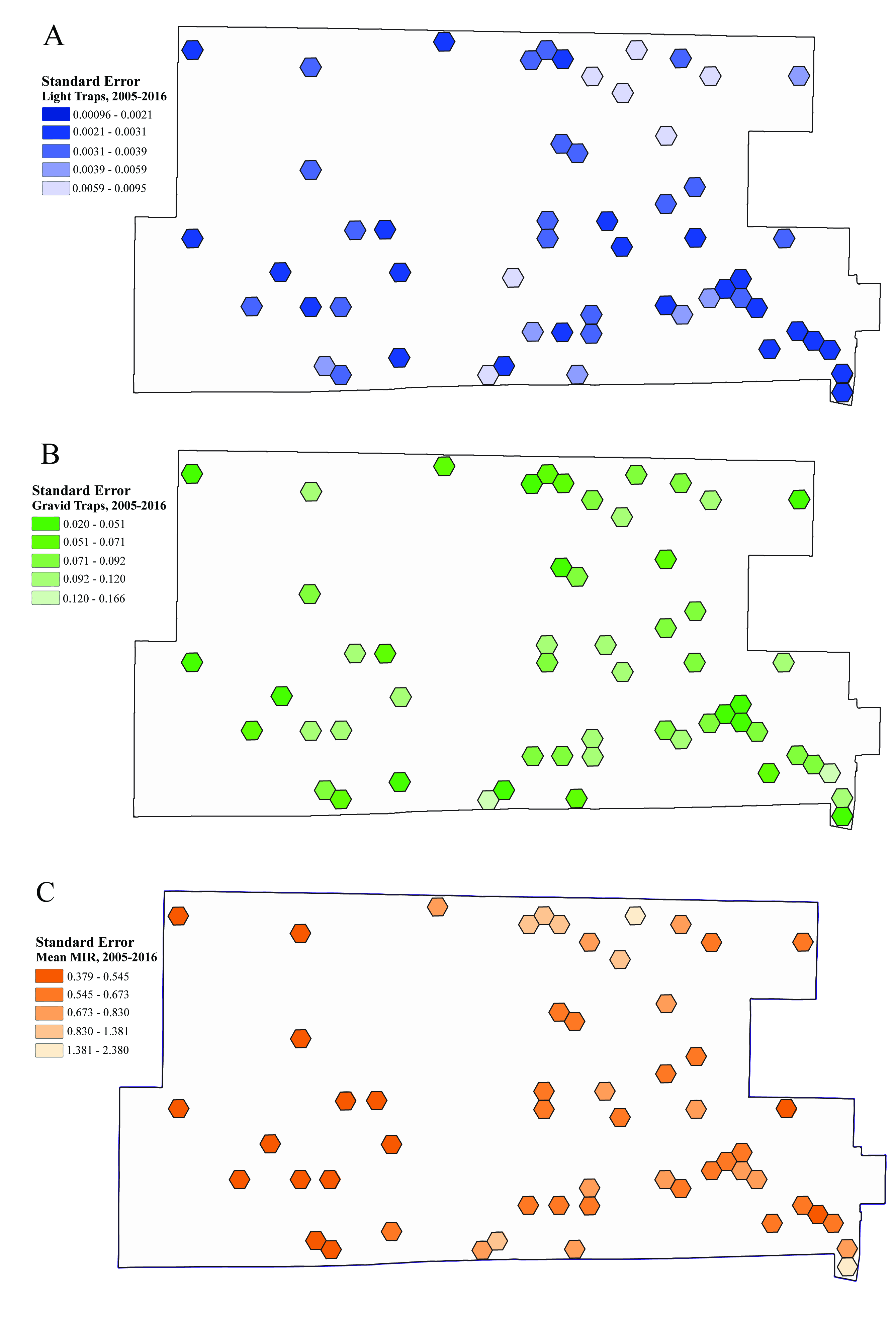

### Supplemental Table 1

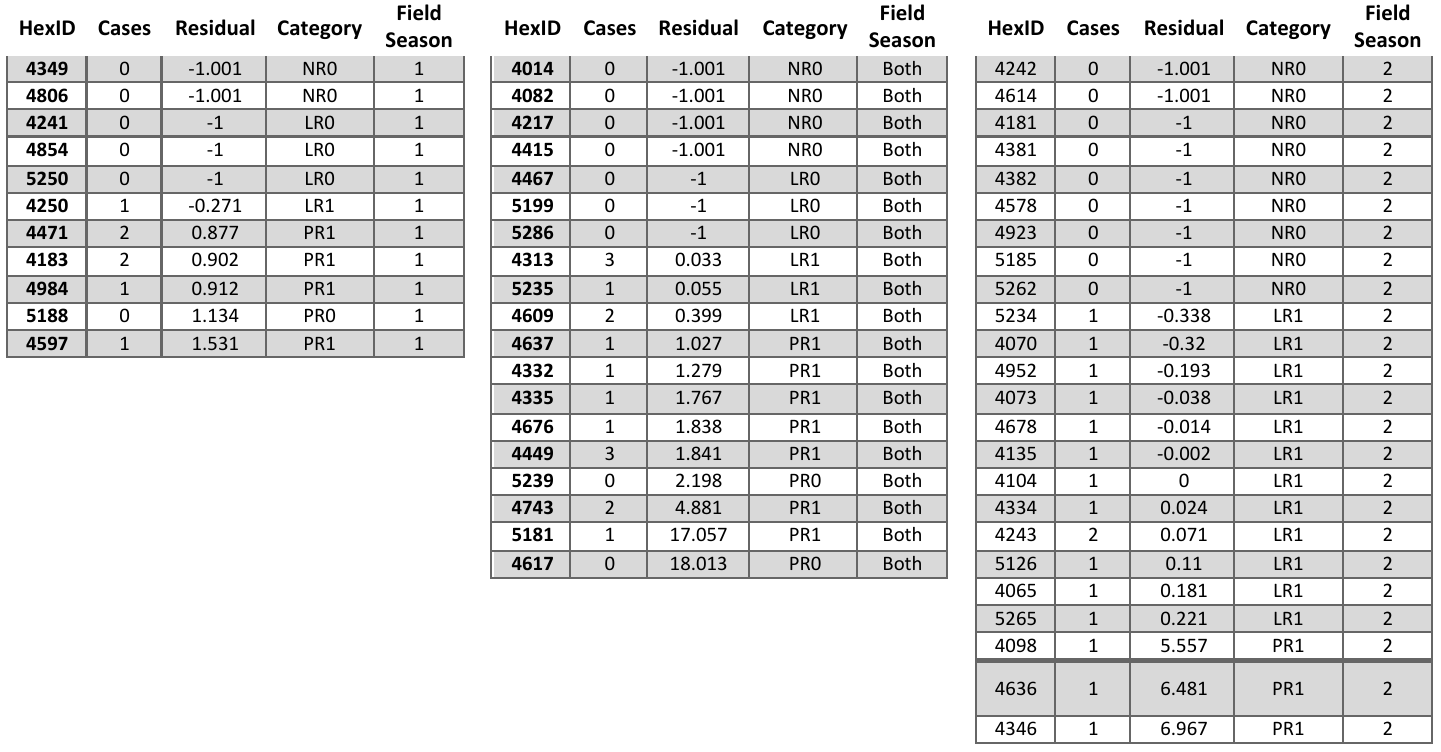

### Supplemental Table 2

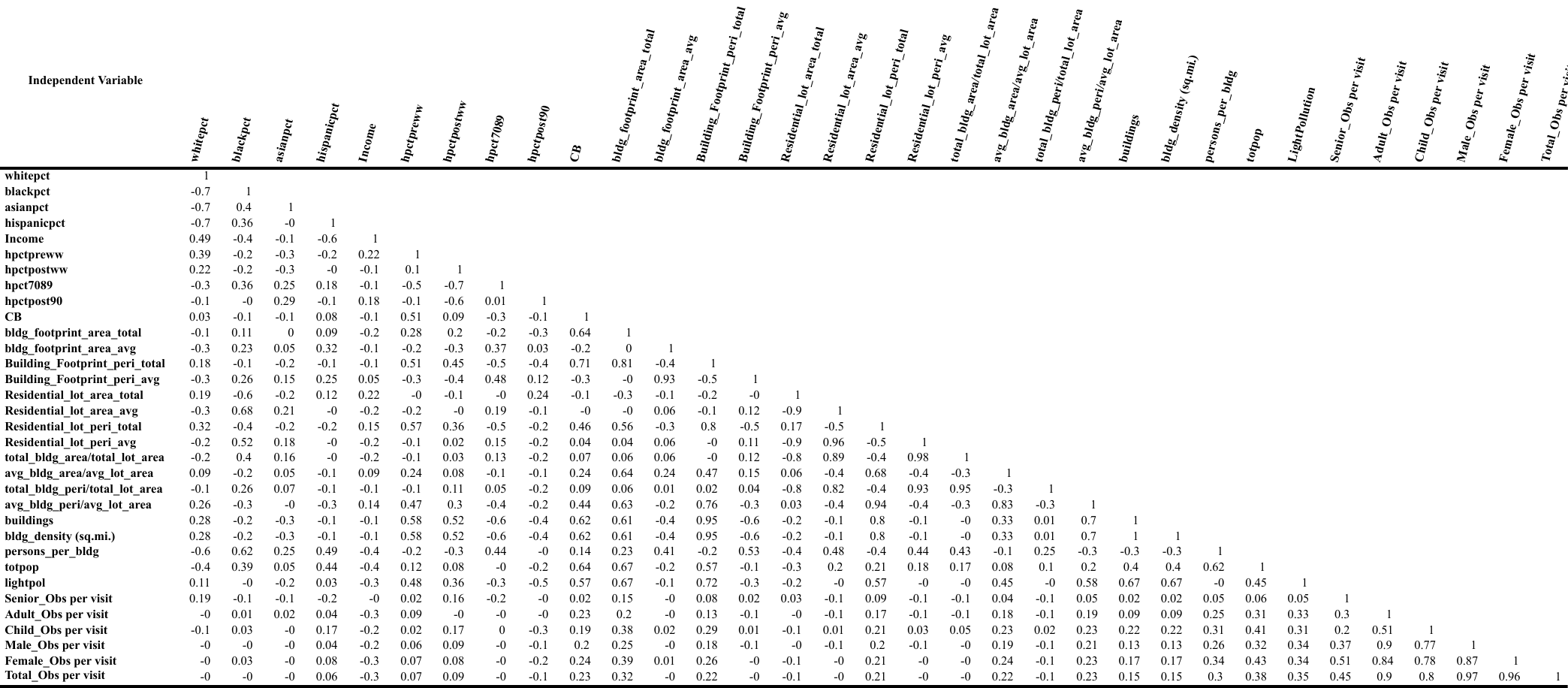

**A.**

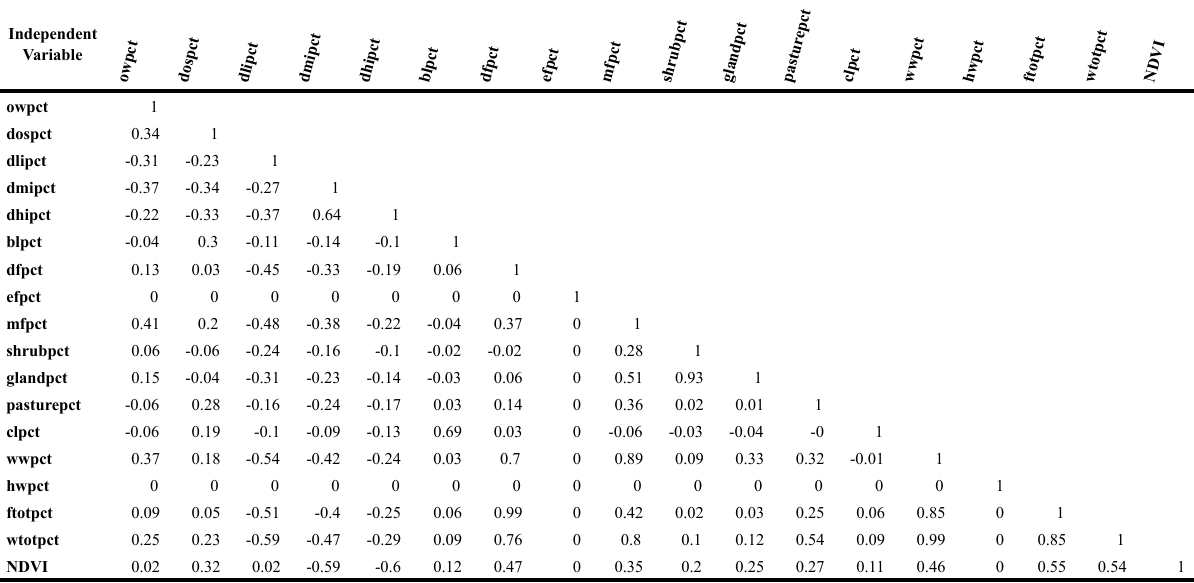

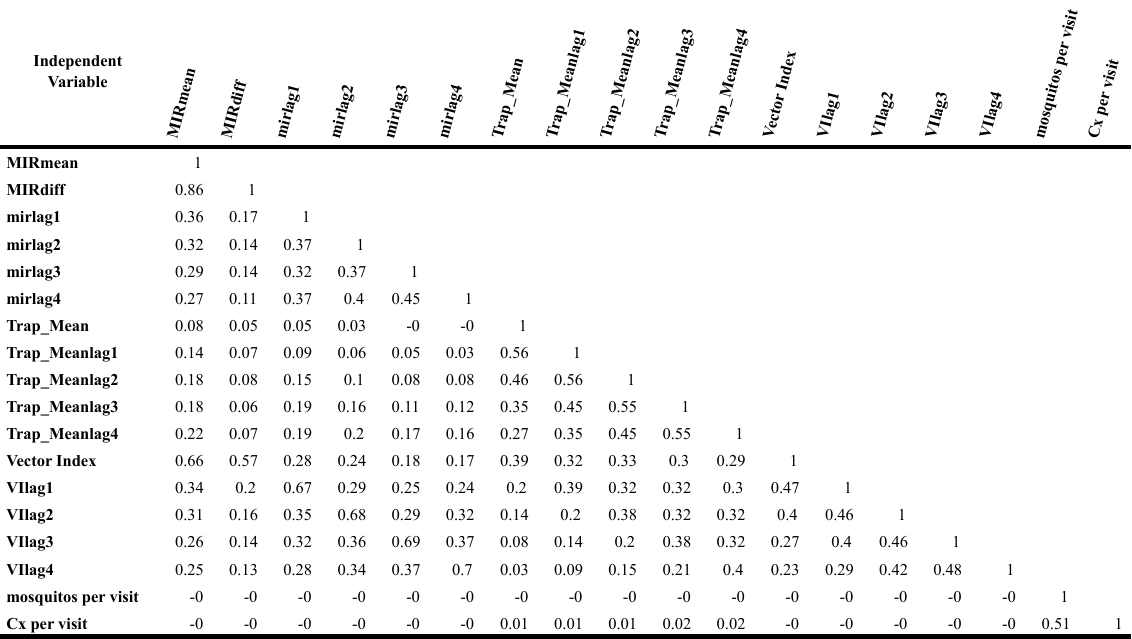

**C.**

**B.**

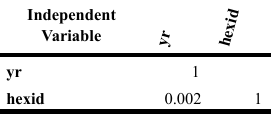

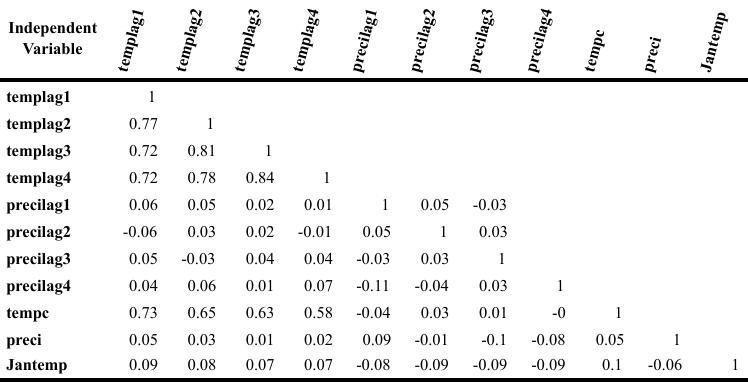

**E.**

**D.**
