## Supplemental Table 3 for "Dynamics of data availability in disease modeling: An example evaluating the trade-offs of ultra-fine-scale factors applied to human West Nile virus disease models in the Chicago area, USA"

| **Covariate** | **Significance Level** | **Data Availability/Work Load to Acquire (Score of 1,2,3,4)** | **Mean Covariate Value** | **HLC/Human Obervation Covariate?** |  | **Covariate** | **Significance Level** | **Data Availability/Work Load to Acquire (Score of 1,2,3,4)** | **Mean Covariate Value** | **HLC/Human Obervation Covariate?** |  | **Covariate** | **Significance Level** | **Data Availability/Work Load to Acquire (Score of 1,2,3,4)** | **Mean Covariate Value** | **HLC/Human Obervation Covariate?** |  | **Covariate** | **Significance Level** | **Data Availability/Work Load to Acquire (Score of 1,2,3,4)** | **Mean Covariate Value** | **HLC/Human Obervation Covariate?** |
| --- | --- | --- | --- | --- | --- | --- | --- | --- | --- | --- | --- | --- | --- | --- | --- | --- | --- | --- | --- | --- | --- | --- |
| preci | 1 | 1 | 1.00 | No |  | asianpct | 3 | 2 | 1.50 | No |  | dhipct | 1 | 2 | 0.50 | No |  | dospct | 1 | 2 | 0.50 | No |
| preci | 1 | 1 | 1.00 | No |  | asianpct | 1 | 2 | 0.50 | No |  | dhipct | 1 | 2 | 0.50 | No |  | dospct | 1 | 2 | 0.50 | No |
| preci | 1 | 1 | 1.00 | No |  | asianpct | 3 | 2 | 1.50 | No |  | dhipct | 1 | 2 | 0.50 | No |  | dospct | 1 | 2 | 0.50 | No |
| preci | 1 | 1 | 1.00 | No |  | asianpct | 1 | 2 | 0.50 | No |  | dhipct | 1 | 2 | 0.50 | No |  | dospct | 1 | 2 | 0.50 | No |
| preci | 1 | 1 | 1.00 | No |  | asianpct | 1 | 2 | 0.50 | No |  | dhipct | 1 | 2 | 0.50 | No |  | dospct | 1 | 2 | 0.50 | No |
| preci | 1 | 1 | 1.00 | No |  | blackpct | 1 | 2 | 0.50 | No |  | dhipct | 1 | 2 | 0.50 | No |  | dospct | 1 | 2 | 0.50 | No |
| preci | 1 | 1 | 1.00 | No |  | blackpct | 1 | 2 | 0.50 | No |  | dhipct | 1 | 2 | 0.50 | No |  | dtotpct | 1 | 2 | 0.50 | No |
| preci | 1 | 1 | 1.00 | No |  | blackpct | 1 | 2 | 0.50 | No |  | dhipct | 1 | 2 | 0.50 | No |  | dtotpct | 1 | 2 | 0.50 | No |
| preci | 2 | 1 | 2.00 | No |  | blackpct | 1 | 2 | 0.50 | No |  | dhipct | 1 | 2 | 0.50 | No |  | dtotpct | 1 | 2 | 0.50 | No |
| preci | 1 | 1 | 1.00 | No |  | blackpct | 1 | 2 | 0.50 | No |  | dhipct | 1 | 2 | 0.50 | No |  | dtotpct | 1 | 2 | 0.50 | No |
| preci | 2 | 1 | 2.00 | No |  | blackpct | 1 | 2 | 0.50 | No |  | dlipct | 1 | 2 | 0.50 | No |  | dtotpct | 1 | 2 | 0.50 | No |
| preci | 1 | 1 | 1.00 | No |  | blackpct | 1 | 2 | 0.50 | No |  | dlipct | 1 | 2 | 0.50 | No |  | ftotpct | 1 | 2 | 0.50 | No |
| preci | 1 | 1 | 1.00 | No |  | blackpct | 1 | 2 | 0.50 | No |  | dlipct | 1 | 2 | 0.50 | No |  | ftotpct | 1 | 2 | 0.50 | No |
| preci | 1 | 1 | 1.00 | No |  | blackpct | 1 | 2 | 0.50 | No |  | dlipct | 1 | 2 | 0.50 | No |  | ftotpct | 1 | 2 | 0.50 | No |
| tempc | 1 | 1 | 1.00 | No |  | blackpct | 1 | 2 | 0.50 | No |  | dlipct | 1 | 2 | 0.50 | No |  | ftotpct | 1 | 2 | 0.50 | No |
| tempc | 1 | 1 | 1.00 | No |  | blackpct | 1 | 2 | 0.50 | No |  | dlipct | 1 | 2 | 0.50 | No |  | ftotpct | 1 | 2 | 0.50 | No |
| tempc | 1 | 1 | 1.00 | No |  | blackpct | 1 | 2 | 0.50 | No |  | dlipct | 1 | 2 | 0.50 | No |  | glandpct | 1 | 2 | 0.50 | No |
| tempc | 1 | 1 | 1.00 | No |  | blackpct | 1 | 2 | 0.50 | No |  | dlipct | 1 | 2 | 0.50 | No |  | glandpct | 1 | 2 | 0.50 | No |
| tempc | 1 | 1 | 1.00 | No |  | blackpct | 1 | 2 | 0.50 | No |  | dlipct | 1 | 2 | 0.50 | No |  | glandpct | 1 | 2 | 0.50 | No |
| tempc | 1 | 1 | 1.00 | No |  | blackpct | 1 | 2 | 0.50 | No |  | dlipct | 1 | 2 | 0.50 | No |  | glandpct | 1 | 2 | 0.50 | No |
| tempc | 1 | 1 | 1.00 | No |  | blackpct | 1 | 2 | 0.50 | No |  | dlipct | 1 | 2 | 0.50 | No |  | glandpct | 1 | 2 | 0.50 | No |
| tempc | 1 | 1 | 1.00 | No |  | blpct | 1 | 2 | 0.50 | No |  | dlipct | 1 | 2 | 0.50 | No |  | glandpct | 1 | 2 | 0.50 | No |
| tempc | 2 | 1 | 2.00 | No |  | blpct | 1 | 2 | 0.50 | No |  | dmipct | 1 | 2 | 0.50 | No |  | glandpct | 1 | 2 | 0.50 | No |
| tempc | 1 | 1 | 1.00 | No |  | blpct | 1 | 2 | 0.50 | No |  | dmipct | 1 | 2 | 0.50 | No |  | glandpct | 1 | 2 | 0.50 | No |
| tempc | 2 | 1 | 2.00 | No |  | blpct | 1 | 2 | 0.50 | No |  | dmipct | 1 | 2 | 0.50 | No |  | glandpct | 1 | 2 | 0.50 | No |
| tempc | 1 | 1 | 1.00 | No |  | blpct | 1 | 2 | 0.50 | No |  | dmipct | 1 | 2 | 0.50 | No |  | glandpct | 1 | 2 | 0.50 | No |
| tempc | 1 | 1 | 1.00 | No |  | blpct | 1 | 2 | 0.50 | No |  | dmipct | 1 | 2 | 0.50 | No |  | glandpct | 1 | 2 | 0.50 | No |
| Yr | 1 | 1 | 1.00 | No |  | blpct | 1 | 2 | 0.50 | No |  | dmipct | 1 | 2 | 0.50 | No |  | hispanicpct | 1 | 2 | 0.50 | No |
| Yr | 1 | 1 | 1.00 | No |  | blpct | 3 | 2 | 1.50 | No |  | dmipct | 1 | 2 | 0.50 | No |  | hispanicpct | 1 | 2 | 0.50 | No |
| Yr | 1 | 1 | 1.00 | No |  | blpct | 3 | 2 | 1.50 | No |  | dmipct | 3 | 2 | 1.50 | No |  | hispanicpct | 1 | 2 | 0.50 | No |
| Yr | 1 | 1 | 1.00 | No |  | blpct | 3 | 2 | 1.50 | No |  | dmipct | 4 | 2 | 2.00 | No |  | hispanicpct | 1 | 2 | 0.50 | No |
| Yr | 1 | 1 | 1.00 | No |  | blpct | 3 | 2 | 1.50 | No |  | dmipct | 1 | 2 | 0.50 | No |  | hispanicpct | 1 | 2 | 0.50 | No |
| Yr | 1 | 1 | 1.00 | No |  | blpct | 3 | 2 | 1.50 | No |  | dmipct | 3 | 2 | 1.50 | No |  | hispanicpct | 1 | 2 | 0.50 | No |
| Yr | 1 | 1 | 1.00 | No |  | ccpct | 1 | 2 | 0.50 | No |  | dmipct | 1 | 2 | 0.50 | No |  | hispanicpct | 1 | 2 | 0.50 | No |
| Yr | 1 | 1 | 1.00 | No |  | ccpct | 1 | 2 | 0.50 | No |  | dmipct | 1 | 2 | 0.50 | No |  | hispanicpct | 1 | 2 | 0.50 | No |
| Yr | 1 | 1 | 1.00 | No |  | ccpct | 1 | 2 | 0.50 | No |  | dmipct | 1 | 2 | 0.50 | No |  | Income | 1 | 2 | 0.50 | No |
| Yr | 1 | 1 | 1.00 | No |  | dfpct | 1 | 2 | 0.50 | No |  | dmipct | 1 | 2 | 0.50 | No |  | Income | 1 | 2 | 0.50 | No |
| Yr | 1 | 1 | 1.00 | No |  | dfpct | 1 | 2 | 0.50 | No |  | dmipct | 1 | 2 | 0.50 | No |  | Income | 1 | 2 | 0.50 | No |
| # bldgs | 1 | 2 | 0.50 | No |  | dfpct | 1 | 2 | 0.50 | No |  | dmipct | 1 | 2 | 0.50 | No |  | Income | 1 | 2 | 0.50 | No |
| # bldgs | 1 | 2 | 0.50 | No |  | dfpct | 1 | 2 | 0.50 | No |  | dmipct | 1 | 2 | 0.50 | No |  | Income | 1 | 2 | 0.50 | No |
| # bldgs | 1 | 2 | 0.50 | No |  | dfpct | 1 | 2 | 0.50 | No |  | dospct | 1 | 2 | 0.50 | No |  | Income | 1 | 2 | 0.50 | No |
| asianpct | 1 | 2 | 0.50 | No |  | dfpct | 1 | 2 | 0.50 | No |  | dospct | 1 | 2 | 0.50 | No |  | Income | 1 | 2 | 0.50 | No |
| asianpct | 1 | 2 | 0.50 | No |  | dfpct | 1 | 2 | 0.50 | No |  | dospct | 1 | 2 | 0.50 | No |  | Income | 1 | 2 | 0.50 | No |
| asianpct | 1 | 2 | 0.50 | No |  | dfpct | 2 | 2 | 1.00 | No |  | dospct | 1 | 2 | 0.50 | No |  | Income | 1 | 2 | 0.50 | No |
| asianpct | 1 | 2 | 0.50 | No |  | dfpct | 2 | 2 | 1.00 | No |  | dospct | 1 | 2 | 0.50 | No |  | Jantemp | 1 | 2 | 0.50 | No |
| asianpct | 1 | 2 | 0.50 | No |  | dfpct | 1 | 2 | 0.50 | No |  | dospct | 1 | 2 | 0.50 | No |  | Jantemp | 1 | 2 | 0.50 | No |
| asianpct | 1 | 2 | 0.50 | No |  | dfpct | 1 | 2 | 0.50 | No |  | dospct | 1 | 2 | 0.50 | No |  | Jantemp | 1 | 2 | 0.50 | No |
| asianpct | 1 | 2 | 0.50 | No |  | dhipct | 1 | 2 | 0.50 | No |  | dospct | 1 | 2 | 0.50 | No |  | Jantemp | 1 | 2 | 0.50 | No |

| **Covariate** | **Significance Level** | **Data Availability/Work Load to Acquire (Score of 1,2,3,4)** | **Mean Covariate Value** | **HLC/Human Obervation Covariate?** |  | **Covariate** | **Significance Level** | **Data Availability/Work Load to Acquire (Score of 1,2,3,4)** | **Mean Covariate Value** | **HLC/Human Obervation Covariate?** |  | **Covariate** | **Significance Level** | **Data Availability/Work Load to Acquire (Score of 1,2,3,4)** | **Mean Covariate Value** | **HLC/Human Obervation Covariate?** |  | **Covariate** | **Significance Level** | **Data Availability/Work Load to Acquire (Score of 1,2,3,4)** | **Mean Covariate Value** | **HLC/Human Obervation Covariate?** |  | **Covariate** | **Significance Level** | **Data Availability/Work Load to Acquire (Score of 1,2,3,4)** | **Mean Covariate Value** | **HLC/Human Obervation Covariate?** |
| --- | --- | --- | --- | --- | --- | --- | --- | --- | --- | --- | --- | --- | --- | --- | --- | --- | --- | --- | --- | --- | --- | --- | --- | --- | --- | --- | --- | --- |
| Jantemp | 1 | 2 | 0.50 | No |  | precilag1 | 1 | 2 | 0.50 | No |  | templag1 | 1 | 2 | 0.50 | No |  | templag4 | 1 | 2 | 0.50 | No |  | wwpct | 1 | 2 | 0.50 | No |
| Jantemp | 1 | 2 | 0.50 | No |  | precilag2 | 1 | 2 | 0.50 | No |  | templag1 | 1 | 2 | 0.50 | No |  | templag4 | 2 | 2 | 1.00 | No |  | wwpct | 1 | 2 | 0.50 | No |
| Jantemp | 1 | 2 | 0.50 | No |  | precilag2 | 1 | 2 | 0.50 | No |  | templag1 | 1 | 2 | 0.50 | No |  | templag4 | 1 | 2 | 0.50 | No |  | wwpct | 1 | 2 | 0.50 | No |
| Jantemp | 1 | 2 | 0.50 | No |  | precilag2 | 1 | 2 | 0.50 | No |  | templag1 | 1 | 2 | 0.50 | No |  | templag4 | 2 | 2 | 1.00 | No |  | wwpct | 1 | 2 | 0.50 | No |
| Jantemp | 1 | 2 | 0.50 | No |  | precilag2 | 1 | 2 | 0.50 | No |  | templag1 | 1 | 2 | 0.50 | No |  | templag4 | 2 | 2 | 1.00 | No |  | wwpct | 1 | 2 | 0.50 | No |
| Jantemp | 1 | 2 | 0.50 | No |  | precilag2 | 1 | 2 | 0.50 | No |  | templag1 | 1 | 2 | 0.50 | No |  | templag4 | 1 | 2 | 0.50 | No |  | abund | 1 | 3 | 0.33 | No |
| Jantemp | 1 | 2 | 0.50 | No |  | precilag2 | 1 | 2 | 0.50 | No |  | templag1 | 1 | 2 | 0.50 | No |  | templag4 | 1 | 2 | 0.50 | No |  | abund | 1 | 3 | 0.33 | No |
| mfpct | 1 | 2 | 0.50 | No |  | precilag2 | 1 | 2 | 0.50 | No |  | templag1 | 1 | 2 | 0.50 | No |  | templag4 | 2 | 2 | 1.00 | No |  | abund | 1 | 3 | 0.33 | No |
| mfpct | 1 | 2 | 0.50 | No |  | precilag2 | 1 | 2 | 0.50 | No |  | templag1 | 1 | 2 | 0.50 | No |  | templag4 | 1 | 2 | 0.50 | No |  | abund | 1 | 3 | 0.33 | No |
| mfpct | 1 | 2 | 0.50 | No |  | precilag2 | 1 | 2 | 0.50 | No |  | templag2 | 1 | 2 | 0.50 | No |  | templag4 | 1 | 2 | 0.50 | No |  | abund | 1 | 3 | 0.33 | No |
| mfpct | 1 | 2 | 0.50 | No |  | precilag2 | 1 | 2 | 0.50 | No |  | templag2 | 1 | 2 | 0.50 | No |  | templag4 | 1 | 2 | 0.50 | No |  | abund | 1 | 3 | 0.33 | No |
| mfpct | 1 | 2 | 0.50 | No |  | precilag3 | 1 | 2 | 0.50 | No |  | templag2 | 1 | 2 | 0.50 | No |  | templag4 | 1 | 2 | 0.50 | No |  | abund | 1 | 3 | 0.33 | No |
| mfpct | 1 | 2 | 0.50 | No |  | precilag3 | 1 | 2 | 0.50 | No |  | templag2 | 1 | 2 | 0.50 | No |  | templag4 | 3 | 2 | 1.50 | No |  | abund | 1 | 3 | 0.33 | No |
| mfpct | 1 | 2 | 0.50 | No |  | precilag3 | 1 | 2 | 0.50 | No |  | templag2 | 1 | 2 | 0.50 | No |  | totpop | 3 | 2 | 1.50 | No |  | abund | 1 | 3 | 0.33 | No |
| mfpct | 1 | 2 | 0.50 | No |  | precilag3 | 1 | 2 | 0.50 | No |  | templag2 | 1 | 2 | 0.50 | No |  | totpop | 2 | 2 | 1.00 | No |  | abund | 1 | 3 | 0.33 | No |
| mfpct | 1 | 2 | 0.50 | No |  | precilag3 | 1 | 2 | 0.50 | No |  | templag2 | 1 | 2 | 0.50 | No |  | totpop | 1 | 2 | 0.50 | No |  | abund | 1 | 3 | 0.33 | No |
| mfpct | 1 | 2 | 0.50 | No |  | precilag3 | 1 | 2 | 0.50 | No |  | templag2 | 1 | 2 | 0.50 | No |  | totpop | 2 | 2 | 1.00 | No |  | abund | 1 | 3 | 0.33 | No |
| owpct | 1 | 2 | 0.50 | No |  | precilag3 | 1 | 2 | 0.50 | No |  | templag2 | 1 | 2 | 0.50 | No |  | totpop | 1 | 2 | 0.50 | No |  | abund | 1 | 3 | 0.33 | No |
| owpct | 1 | 2 | 0.50 | No |  | precilag3 | 1 | 2 | 0.50 | No |  | templag2 | 1 | 2 | 0.50 | No |  | totpop | 2 | 2 | 1.00 | No |  | abund | 1 | 3 | 0.33 | No |
| owpct | 1 | 2 | 0.50 | No |  | precilag3 | 1 | 2 | 0.50 | No |  | templag2 | 1 | 2 | 0.50 | No |  | totpop | 1 | 2 | 0.50 | No |  | abund | 1 | 3 | 0.33 | No |
| owpct | 1 | 2 | 0.50 | No |  | precilag3 | 1 | 2 | 0.50 | No |  | templag2 | 1 | 2 | 0.50 | No |  | totpop | 1 | 2 | 0.50 | No |  | abundlag1 | 1 | 3 | 0.33 | No |
| owpct | 1 | 2 | 0.50 | No |  | precilag4 | 2 | 2 | 1.00 | No |  | templag2 | 1 | 2 | 0.50 | No |  | totpop | 1 | 2 | 0.50 | No |  | abundlag1 | 1 | 3 | 0.33 | No |
| owpct | 1 | 2 | 0.50 | No |  | precilag4 | 2 | 2 | 1.00 | No |  | templag2 | 1 | 2 | 0.50 | No |  | totpop | 1 | 2 | 0.50 | No |  | abundlag1 | 1 | 3 | 0.33 | No |
| owpct | 1 | 2 | 0.50 | No |  | precilag4 | 2 | 2 | 1.00 | No |  | templag2 | 1 | 2 | 0.50 | No |  | totpop | 1 | 2 | 0.50 | No |  | abundlag1 | 1 | 3 | 0.33 | No |
| owpct | 2 | 2 | 1.00 | No |  | precilag4 | 1 | 2 | 0.50 | No |  | templag2 | 1 | 2 | 0.50 | No |  | totpop | 1 | 2 | 0.50 | No |  | abundlag1 | 1 | 3 | 0.33 | No |
| owpct | 2 | 2 | 1.00 | No |  | precilag4 | 2 | 2 | 1.00 | No |  | templag2 | 1 | 2 | 0.50 | No |  | totpop | 1 | 2 | 0.50 | No |  | abundlag1 | 1 | 3 | 0.33 | No |
| owpct | 1 | 2 | 0.50 | No |  | precilag4 | 1 | 2 | 0.50 | No |  | templag2 | 1 | 2 | 0.50 | No |  | totpop | 1 | 2 | 0.50 | No |  | abundlag1 | 2 | 3 | 0.67 | No |
| owpct | 1 | 2 | 0.50 | No |  | precilag4 | 1 | 2 | 0.50 | No |  | templag2 | 1 | 2 | 0.50 | No |  | totpop | 1 | 2 | 0.50 | No |  | abundlag1 | 1 | 3 | 0.33 | No |
| pasturepct | 1 | 2 | 0.50 | No |  | precilag4 | 1 | 2 | 0.50 | No |  | templag3 | 2 | 2 | 1.00 | No |  | totpop | 2 | 2 | 1.00 | No |  | abundlag1 | 1 | 3 | 0.33 | No |
| pasturepct | 1 | 2 | 0.50 | No |  | precilag4 | 1 | 2 | 0.50 | No |  | templag3 | 3 | 2 | 1.50 | No |  | totpop | 1 | 2 | 0.50 | No |  | abundlag1 | 1 | 3 | 0.33 | No |
| pasturepct | 1 | 2 | 0.50 | No |  | precilag4 | 1 | 2 | 0.50 | No |  | templag3 | 2 | 2 | 1.00 | No |  | totpop | 1 | 2 | 0.50 | No |  | abundlag1 | 1 | 3 | 0.33 | No |
| pasturepct | 1 | 2 | 0.50 | No |  | precilag4 | 1 | 2 | 0.50 | No |  | templag3 | 2 | 2 | 1.00 | No |  | totpop | 1 | 2 | 0.50 | No |  | abundlag1 | 1 | 3 | 0.33 | No |
| pasturepct | 1 | 2 | 0.50 | No |  | precilag4 | 1 | 2 | 0.50 | No |  | templag3 | 2 | 2 | 1.00 | No |  | totpop | 2 | 2 | 1.00 | No |  | abundlag1 | 1 | 3 | 0.33 | No |
| pasturepct | 1 | 2 | 0.50 | No |  | precilag4 | 2 | 2 | 1.00 | No |  | templag3 | 1 | 2 | 0.50 | No |  | whitepct | 1 | 2 | 0.50 | No |  | abundlag1 | 1 | 3 | 0.33 | No |
| pasturepct | 2 | 2 | 1.00 | No |  | precilag4 | 1 | 2 | 0.50 | No |  | templag3 | 3 | 2 | 1.50 | No |  | whitepct | 1 | 2 | 0.50 | No |  | abundlag2 | 1 | 3 | 0.33 | No |
| precilag1 | 2 | 2 | 1.00 | No |  | precilag4 | 2 | 2 | 1.00 | No |  | templag3 | 1 | 2 | 0.50 | No |  | whitepct | 1 | 2 | 0.50 | No |  | abundlag2 | 1 | 3 | 0.33 | No |
| precilag1 | 2 | 2 | 1.00 | No |  | precilag4 | 1 | 2 | 0.50 | No |  | templag3 | 3 | 2 | 1.50 | No |  | whitepct | 1 | 2 | 0.50 | No |  | abundlag2 | 1 | 3 | 0.33 | No |
| precilag1 | 2 | 2 | 1.00 | No |  | precilag4 | 2 | 2 | 1.00 | No |  | templag3 | 3 | 2 | 1.50 | No |  | whitepct | 1 | 2 | 0.50 | No |  | abundlag2 | 1 | 3 | 0.33 | No |
| precilag1 | 2 | 2 | 1.00 | No |  | precilag4 | 1 | 2 | 0.50 | No |  | templag3 | 1 | 2 | 0.50 | No |  | whitepct | 1 | 2 | 0.50 | No |  | abundlag2 | 1 | 3 | 0.33 | No |
| precilag1 | 2 | 2 | 1.00 | No |  | precilag4 | 1 | 2 | 0.50 | No |  | templag3 | 3 | 2 | 1.50 | No |  | whitepct | 1 | 2 | 0.50 | No |  | abundlag2 | 1 | 3 | 0.33 | No |
| precilag1 | 2 | 2 | 1.00 | No |  | precilag4 | 1 | 2 | 0.50 | No |  | templag3 | 1 | 2 | 0.50 | No |  | whitepct | 1 | 2 | 0.50 | No |  | abundlag2 | 1 | 3 | 0.33 | No |
| precilag1 | 2 | 2 | 1.00 | No |  | shrubpct | 1 | 2 | 0.50 | No |  | templag3 | 3 | 2 | 1.50 | No |  | whitepct | 1 | 2 | 0.50 | No |  | abundlag2 | 1 | 3 | 0.33 | No |
| precilag1 | 2 | 2 | 1.00 | No |  | shrubpct | 1 | 2 | 0.50 | No |  | templag3 | 2 | 2 | 1.00 | No |  | whitepct | 1 | 2 | 0.50 | No |  | abundlag2 | 1 | 3 | 0.33 | No |
| precilag1 | 1 | 2 | 0.50 | No |  | shrubpct | 1 | 2 | 0.50 | No |  | templag3 | 2 | 2 | 1.00 | No |  | whitepct | 1 | 2 | 0.50 | No |  | abundlag2 | 1 | 3 | 0.33 | No |
| precilag1 | 2 | 2 | 1.00 | No |  | shrubpct | 1 | 2 | 0.50 | No |  | templag3 | 2 | 2 | 1.00 | No |  | whitepct | 1 | 2 | 0.50 | No |  | abundlag2 | 1 | 3 | 0.33 | No |
| precilag1 | 2 | 2 | 1.00 | No |  | shrubpct | 1 | 2 | 0.50 | No |  | templag3 | 2 | 2 | 1.00 | No |  | wtotpct | 1 | 2 | 0.50 | No |  | abundlag2 | 1 | 3 | 0.33 | No |
| precilag1 | 2 | 2 | 1.00 | No |  | shrubpct | 1 | 2 | 0.50 | No |  | templag3 | 1 | 2 | 0.50 | No |  | wtotpct | 1 | 2 | 0.50 | No |  | abundlag2 | 1 | 3 | 0.33 | No |
| precilag1 | 2 | 2 | 1.00 | No |  | shrubpct | 1 | 2 | 0.50 | No |  | templag3 | 1 | 2 | 0.50 | No |  | wtotpct | 1 | 2 | 0.50 | No |  | abundlag3 | 1 | 3 | 0.33 | No |
| precilag1 | 3 | 2 | 1.50 | No |  | templag1 | 2 | 2 | 1.00 | No |  | templag3 | 2 | 2 | 1.00 | No |  | wtotpct | 1 | 2 | 0.50 | No |  | abundlag3 | 1 | 3 | 0.33 | No |
| precilag1 | 1 | 2 | 0.50 | No |  | templag1 | 1 | 2 | 0.50 | No |  | templag3 | 1 | 2 | 0.50 | No |  | wtotpct | 1 | 2 | 0.50 | No |  | abundlag3 | 1 | 3 | 0.33 | No |
| precilag1 | 2 | 2 | 1.00 | No |  | templag1 | 1 | 2 | 0.50 | No |  | templag3 | 1 | 2 | 0.50 | No |  | wwpct | 1 | 2 | 0.50 | No |  | abundlag3 | 1 | 3 | 0.33 | No |
| precilag1 | 1 | 2 | 0.50 | No |  | templag1 | 1 | 2 | 0.50 | No |  | templag3 | 1 | 2 | 0.50 | No |  | wwpct | 1 | 2 | 0.50 | No |  | abundlag3 | 1 | 3 | 0.33 | No |
| precilag1 | 3 | 2 | 1.50 | No |  | templag1 | 2 | 2 | 1.00 | No |  | templag3 | 1 | 2 | 0.50 | No |  | wwpct | 1 | 2 | 0.50 | No |  | abundlag3 | 1 | 3 | 0.33 | No |
| precilag1 | 1 | 2 | 0.50 | No |  | templag1 | 1 | 2 | 0.50 | No |  | templag3 | 2 | 2 | 1.00 | No |  | wwpct | 1 | 2 | 0.50 | No |  | abundlag3 | 1 | 3 | 0.33 | No |
| precilag1 | 1 | 2 | 0.50 | No |  | templag1 | 1 | 2 | 0.50 | No |  | templag4 | 2 | 2 | 1.00 | No |  | wwpct | 1 | 2 | 0.50 | No |  | abundlag3 | 1 | 3 | 0.33 | No |

| **Covariate** | **Significance Level** | **Data Availability/Work Load to Acquire (Score of 1,2,3,4)** | **Mean Covariate Value** | **HLC/Human Obervation Covariate?** |  | **Covariate** | **Significance Level** | **Data Availability/Work Load to Acquire (Score of 1,2,3,4)** | **Mean Covariate Value** | **HLC/Human Obervation Covariate?** |  | **Covariate** | **Significance Level** | **Data Availability/Work Load to Acquire (Score of 1,2,3,4)** | **Mean Covariate Value** | **HLC/Human Obervation Covariate?** |  | **Covariate** | **Significance Level** | **Data Availability/Work Load to Acquire (Score of 1,2,3,4)** | **Mean Covariate Value** | **HLC/Human Obervation Covariate?** |
| --- | --- | --- | --- | --- | --- | --- | --- | --- | --- | --- | --- | --- | --- | --- | --- | --- | --- | --- | --- | --- | --- | --- |
| abundlag3 | 1 | 3 | 0.33 | No |  | hpctpostww | 1 | 3 | 0.33 | No |  | mirlag2 | 1 | 3 | 0.33 | No |  | mirlag4 | 2 | 3 | 0.67 | No |
| abundlag3 | 1 | 3 | 0.33 | No |  | hpctpostww | 1 | 3 | 0.33 | No |  | mirlag2 | 1 | 3 | 0.33 | No |  | mirlag4 | 2 | 3 | 0.67 | No |
| abundlag3 | 1 | 3 | 0.33 | No |  | hpctpostww | 1 | 3 | 0.33 | No |  | mirlag2 | 1 | 3 | 0.33 | No |  | mirlag4 | 1 | 3 | 0.33 | No |
| abundlag3 | 1 | 3 | 0.33 | No |  | hpctpreww | 1 | 3 | 0.33 | No |  | mirlag2 | 1 | 3 | 0.33 | No |  | mirlag4 | 2 | 3 | 0.67 | No |
| abundlag3 | 1 | 3 | 0.33 | No |  | hpctpreww | 1 | 3 | 0.33 | No |  | mirlag2 | 1 | 3 | 0.33 | No |  | mirlag4 | 1 | 3 | 0.33 | No |
| abundlag4 | 1 | 3 | 0.33 | No |  | hpctpreww | 1 | 3 | 0.33 | No |  | mirlag2 | 1 | 3 | 0.33 | No |  | mirlag4 | 2 | 3 | 0.67 | No |
| abundlag4 | 1 | 3 | 0.33 | No |  | hpctpreww | 1 | 3 | 0.33 | No |  | mirlag2 | 1 | 3 | 0.33 | No |  | mirlag4 | 2 | 3 | 0.67 | No |
| abundlag4 | 1 | 3 | 0.33 | No |  | hpctpreww | 1 | 3 | 0.33 | No |  | mirlag2 | 1 | 3 | 0.33 | No |  | mirlag4 | 2 | 3 | 0.67 | No |
| abundlag4 | 1 | 3 | 0.33 | No |  | hpctpreww | 1 | 3 | 0.33 | No |  | mirlag2 | 1 | 3 | 0.33 | No |  | mirlag4 | 1 | 3 | 0.33 | No |
| abundlag4 | 1 | 3 | 0.33 | No |  | hpctpreww | 1 | 3 | 0.33 | No |  | mirlag2 | 1 | 3 | 0.33 | No |  | MIRmean | 1 | 3 | 0.33 | No |
| abundlag4 | 2 | 3 | 0.67 | No |  | hpctpreww | 1 | 3 | 0.33 | No |  | mirlag2 | 1 | 3 | 0.33 | No |  | MIRmean | 1 | 3 | 0.33 | No |
| abundlag4 | 2 | 3 | 0.67 | No |  | hpctpreww | 1 | 3 | 0.33 | No |  | mirlag2 | 1 | 3 | 0.33 | No |  | MIRmean | 1 | 3 | 0.33 | No |
| abundlag4 | 2 | 3 | 0.67 | No |  | hpctpreww | 1 | 3 | 0.33 | No |  | mirlag2 | 1 | 3 | 0.33 | No |  | MIRmean | 1 | 3 | 0.33 | No |
| abundlag4 | 1 | 3 | 0.33 | No |  | hpctpreww | 1 | 3 | 0.33 | No |  | mirlag2 | 1 | 3 | 0.33 | No |  | MIRmean | 1 | 3 | 0.33 | No |
| abundlag4 | 2 | 3 | 0.67 | No |  | Light pol | 1 | 3 | 0.33 | No |  | mirlag2 | 1 | 3 | 0.33 | No |  | MIRmean | 2 | 3 | 0.67 | No |
| abundlag4 | 1 | 3 | 0.33 | No |  | Light pol | 1 | 3 | 0.33 | No |  | mirlag2 | 1 | 3 | 0.33 | No |  | MIRmean | 1 | 3 | 0.33 | No |
| abundlag4 | 1 | 3 | 0.33 | No |  | Light pol | 1 | 3 | 0.33 | No |  | mirlag3 | 2 | 3 | 0.67 | No |  | MIRmean | 1 | 3 | 0.33 | No |
| abundlag4 | 1 | 3 | 0.33 | No |  | Light pol | 1 | 3 | 0.33 | No |  | mirlag3 | 1 | 3 | 0.33 | No |  | MIRmean | 1 | 3 | 0.33 | No |
| CB | 1 | 3 | 0.33 | No |  | Light pol | 1 | 3 | 0.33 | No |  | mirlag3 | 2 | 3 | 0.67 | No |  | MIRmean | 1 | 3 | 0.33 | No |
| CB | 1 | 3 | 0.33 | No |  | Light pol | 1 | 3 | 0.33 | No |  | mirlag3 | 2 | 3 | 0.67 | No |  | MIRmean | 1 | 3 | 0.33 | No |
| CB | 1 | 3 | 0.33 | No |  | MIRdiff | 1 | 3 | 0.33 | No |  | mirlag3 | 2 | 3 | 0.67 | No |  | MIRmean | 1 | 3 | 0.33 | No |
| CB | 1 | 3 | 0.33 | No |  | MIRdiff | 1 | 3 | 0.33 | No |  | mirlag3 | 1 | 3 | 0.33 | No |  | MIRmean | 1 | 3 | 0.33 | No |
| CB | 1 | 3 | 0.33 | No |  | MIRdiff | 1 | 3 | 0.33 | No |  | mirlag3 | 1 | 3 | 0.33 | No |  | resi lot area avg | 1 | 3 | 0.33 | No |
| CB | 1 | 3 | 0.33 | No |  | MIRdiff | 1 | 3 | 0.33 | No |  | mirlag3 | 3 | 3 | 1.00 | No |  | resi lot area avg | 1 | 3 | 0.33 | No |
| hpct7089 | 1 | 3 | 0.33 | No |  | MIRdiff | 1 | 3 | 0.33 | No |  | mirlag3 | 3 | 3 | 1.00 | No |  | resi lot area avg | 1 | 3 | 0.33 | No |
| hpct7089 | 1 | 3 | 0.33 | No |  | MIRdiff | 1 | 3 | 0.33 | No |  | mirlag3 | 2 | 3 | 0.67 | No |  | resi lot area avg | 1 | 3 | 0.33 | No |
| hpct7089 | 1 | 3 | 0.33 | No |  | MIRdiff | 1 | 3 | 0.33 | No |  | mirlag3 | 2 | 3 | 0.67 | No |  | resi lot area total | 1 | 3 | 0.33 | No |
| hpct7089 | 1 | 3 | 0.33 | No |  | MIRdiff | 1 | 3 | 0.33 | No |  | mirlag3 | 2 | 3 | 0.67 | No |  | resi lot area total | 1 | 3 | 0.33 | No |
| hpct7089 | 1 | 3 | 0.33 | No |  | MIRdiff | 1 | 3 | 0.33 | No |  | mirlag3 | 2 | 3 | 0.67 | No |  | resi lot area total | 1 | 3 | 0.33 | No |
| hpct7089 | 1 | 3 | 0.33 | No |  | MIRdiff | 1 | 3 | 0.33 | No |  | mirlag3 | 3 | 3 | 1.00 | No |  | resi lot area total | 1 | 3 | 0.33 | No |
| hpct7089 | 1 | 3 | 0.33 | No |  | MIRdiff | 1 | 3 | 0.33 | No |  | mirlag3 | 3 | 3 | 1.00 | No |  | resi lot area total | 1 | 3 | 0.33 | No |
| hpct7089 | 1 | 3 | 0.33 | No |  | MIRdiff | 1 | 3 | 0.33 | No |  | mirlag3 | 3 | 3 | 1.00 | No |  | resi lot peri avg | 1 | 3 | 0.33 | No |
| hpct7089 | 1 | 3 | 0.33 | No |  | MIRdiff | 1 | 3 | 0.33 | No |  | mirlag3 | 1 | 3 | 0.33 | No |  | resi lot peri avg | 1 | 3 | 0.33 | No |
| hpct7089 | 1 | 3 | 0.33 | No |  | mirlag1 | 1 | 3 | 0.33 | No |  | mirlag3 | 1 | 3 | 0.33 | No |  | resi lot peri avg | 1 | 3 | 0.33 | No |
| hpct7089 | 1 | 3 | 0.33 | No |  | mirlag1 | 1 | 3 | 0.33 | No |  | mirlag3 | 1 | 3 | 0.33 | No |  | resi lot peri avg | 1 | 3 | 0.33 | No |
| hpct7089 | 1 | 3 | 0.33 | No |  | mirlag1 | 1 | 3 | 0.33 | No |  | mirlag3 | 3 | 3 | 1.00 | No |  | resi lot peri avg | 1 | 3 | 0.33 | No |
| hpct7089 | 1 | 3 | 0.33 | No |  | mirlag1 | 1 | 3 | 0.33 | No |  | mirlag3 | 2 | 3 | 0.67 | No |  | resi lot peri avg | 1 | 3 | 0.33 | No |
| hpct7089 | 1 | 3 | 0.33 | No |  | mirlag1 | 1 | 3 | 0.33 | No |  | mirlag3 | 1 | 3 | 0.33 | No |  | resi lot peri avg | 1 | 3 | 0.33 | No |
| hpctpost90 | 1 | 3 | 0.33 | No |  | mirlag1 | 1 | 3 | 0.33 | No |  | mirlag3 | 3 | 3 | 1.00 | No |  | resi lot peri total | 1 | 3 | 0.33 | No |
| hpctpost90 | 2 | 3 | 0.67 | No |  | mirlag1 | 1 | 3 | 0.33 | No |  | mirlag3 | 1 | 3 | 0.33 | No |  | resi lot peri total | 1 | 3 | 0.33 | No |
| hpctpost90 | 1 | 3 | 0.33 | No |  | mirlag1 | 1 | 3 | 0.33 | No |  | mirlag4 | 1 | 3 | 0.33 | No |  | resi lot peri total | 1 | 3 | 0.33 | No |
| hpctpost90 | 1 | 3 | 0.33 | No |  | mirlag1 | 1 | 3 | 0.33 | No |  | mirlag4 | 3 | 3 | 1.00 | No |  | resi lot peri total | 1 | 3 | 0.33 | No |
| hpctpost90 | 1 | 3 | 0.33 | No |  | mirlag1 | 1 | 3 | 0.33 | No |  | mirlag4 | 1 | 3 | 0.33 | No |  | resi lot peri total | 1 | 3 | 0.33 | No |
| hpctpost90 | 1 | 3 | 0.33 | No |  | mirlag1 | 1 | 3 | 0.33 | No |  | mirlag4 | 2 | 3 | 0.67 | No |  | VI | 1 | 3 | 0.33 | No |
| hpctpost90 | 1 | 3 | 0.33 | No |  | mirlag1 | 1 | 3 | 0.33 | No |  | mirlag4 | 1 | 3 | 0.33 | No |  | VI | 1 | 3 | 0.33 | No |
| hpctpost90 | 1 | 3 | 0.33 | No |  | mirlag1 | 1 | 3 | 0.33 | No |  | mirlag4 | 3 | 3 | 1.00 | No |  | VI | 1 | 3 | 0.33 | No |
| hpctpost90 | 1 | 3 | 0.33 | No |  | mirlag1 | 1 | 3 | 0.33 | No |  | mirlag4 | 3 | 3 | 1.00 | No |  | VI | 1 | 3 | 0.33 | No |
| hpctpost90 | 1 | 3 | 0.33 | No |  | mirlag1 | 1 | 3 | 0.33 | No |  | mirlag4 | 3 | 3 | 1.00 | No |  | VI | 1 | 3 | 0.33 | No |
| hpctpost90 | 1 | 3 | 0.33 | No |  | mirlag1 | 1 | 3 | 0.33 | No |  | mirlag4 | 3 | 3 | 1.00 | No |  | VI | 1 | 3 | 0.33 | No |
| hpctpostww | 1 | 3 | 0.33 | No |  | mirlag1 | 1 | 3 | 0.33 | No |  | mirlag4 | 3 | 3 | 1.00 | No |  | VI | 1 | 3 | 0.33 | No |
| hpctpostww | 1 | 3 | 0.33 | No |  | mirlag1 | 1 | 3 | 0.33 | No |  | mirlag4 | 1 | 3 | 0.33 | No |  | VI | 1 | 3 | 0.33 | No |
| hpctpostww | 1 | 3 | 0.33 | No |  | mirlag1 | 1 | 3 | 0.33 | No |  | mirlag4 | 1 | 3 | 0.33 | No |  | VI | 1 | 3 | 0.33 | No |
| hpctpostww | 1 | 3 | 0.33 | No |  | mirlag2 | 1 | 3 | 0.33 | No |  | mirlag4 | 1 | 3 | 0.33 | No |  | VI | 1 | 3 | 0.33 | No |
| hpctpostww | 1 | 3 | 0.33 | No |  | mirlag2 | 1 | 3 | 0.33 | No |  | mirlag4 | 2 | 3 | 0.67 | No |  | VI | 1 | 3 | 0.33 | No |
| hpctpostww | 1 | 3 | 0.33 | No |  | mirlag2 | 1 | 3 | 0.33 | No |  | mirlag4 | 2 | 3 | 0.67 | No |  | VI | 1 | 3 | 0.33 | No |

| **Covariate** | **Significance Level** | **Data Availability/Work Load to Acquire (Score of 1,2,3,4)** | **Mean Covariate Value** | **HLC/Human Obervation Covariate?** |  | **Covariate** | **Significance Level** | **Data Availability/Work Load to Acquire (Score of 1,2,3,4)** | **Mean Covariate Value** | **HLC/Human Obervation Covariate?** |  | **Covariate** | **Significance Level** | **Data Availability/Work Load to Acquire (Score of 1,2,3,4)** | **Mean Covariate Value** | **HLC/Human Obervation Covariate?** |  | **Covariate** | **Significance Level** | **Data Availability/Work Load to Acquire (Score of 1,2,3,4)** | **Mean Covariate Value** | **HLC/Human Obervation Covariate?** |
| --- | --- | --- | --- | --- | --- | --- | --- | --- | --- | --- | --- | --- | --- | --- | --- | --- | --- | --- | --- | --- | --- | --- |
| VIlag1 | 1 | 3 | 0.33 | No |  | VIlag4 | 1 | 3 | 0.33 | No |  | bldg footprint area avg | 1 | 4 | 0.25 | No |  | female obs per visit | 1 | 4 | 0.25 | Yes |
| VIlag1 | 1 | 3 | 0.33 | No |  | VIlag4 | 4 | 3 | 1.33 | No |  | bldg footprint area avg | 1 | 4 | 0.25 | No |  | female obs per visit | 1 | 4 | 0.25 | Yes |
| VIlag1 | 1 | 3 | 0.33 | No |  | VIlag4 | 4 | 3 | 1.33 | No |  | bldg footprint area avg | 1 | 4 | 0.25 | No |  | female obs per visit | 1 | 4 | 0.25 | Yes |
| VIlag1 | 1 | 3 | 0.33 | No |  | Vilag4 | 4 | 3 | 1.33 | No |  | bldg footprint area avg | 1 | 4 | 0.25 | No |  | female obs per visit | 1 | 4 | 0.25 | Yes |
| VIlag1 | 1 | 3 | 0.33 | No |  | VIlag4 | 4 | 3 | 1.33 | No |  | bldg footprint area avg | 4 | 4 | 1.00 | No |  | male obs per visit | 1 | 4 | 0.25 | Yes |
| VIlag1 | 1 | 3 | 0.33 | No |  | VIlag4 | 2 | 3 | 0.67 | No |  | bldg footprint area total | 1 | 4 | 0.25 | No |  | male obs per visit | 1 | 4 | 0.25 | Yes |
| VIlag1 | 1 | 3 | 0.33 | No |  | VIlag4 | 1 | 3 | 0.33 | No |  | bldg footprint area total | 1 | 4 | 0.25 | No |  | male obs per visit | 1 | 4 | 0.25 | Yes |
| VIlag1 | 1 | 3 | 0.33 | No |  | VIlag4 | 4 | 3 | 1.33 | No |  | bldg footprint area total | 1 | 4 | 0.25 | No |  | male obs per visit | 1 | 4 | 0.25 | Yes |
| VIlag1 | 1 | 3 | 0.33 | No |  | VIlag4 | 4 | 3 | 1.33 | No |  | bldg footprint area total | 1 | 4 | 0.25 | No |  | male obs per visit | 1 | 4 | 0.25 | Yes |
| VIlag1 | 1 | 3 | 0.33 | No |  | VIlag4 | 4 | 3 | 1.33 | No |  | bldg footprint area total | 1 | 4 | 0.25 | No |  | male obs per visit | 1 | 4 | 0.25 | Yes |
| VIlag1 | 1 | 3 | 0.33 | No |  | VIlag4 | 1 | 3 | 0.33 | No |  | bldg footprint area total | 1 | 4 | 0.25 | No |  | male obs per visit | 1 | 4 | 0.25 | Yes |
| VIlag1 | 1 | 3 | 0.33 | No |  | VIlag4 | 1 | 3 | 0.33 | No |  | bldg footprint area total | 1 | 4 | 0.25 | No |  | male obs per visit | 1 | 4 | 0.25 | Yes |
| VIlag1 | 1 | 3 | 0.33 | No |  | VIlag4 | 1 | 3 | 0.33 | No |  | bldg footprint peri avg | 1 | 4 | 0.25 | No |  | male obs per visit | 1 | 4 | 0.25 | Yes |
| VIlag2 | 1 | 3 | 0.33 | No |  | VIlag4 | 1 | 3 | 0.33 | No |  | bldg footprint peri avg | 1 | 4 | 0.25 | No |  | mosquitoes per visit | 1 | 4 | 0.25 | Yes |
| VIlag2 | 1 | 3 | 0.33 | No |  | VIlag4 | 1 | 3 | 0.33 | No |  | bldg footprint peri avg | 1 | 4 | 0.25 | No |  | mosquitoes per visit | 1 | 4 | 0.25 | Yes |
| VIlag2 | 1 | 3 | 0.33 | No |  | VIlag4 | 4 | 3 | 1.33 | No |  | bldg footprint peri avg | 1 | 4 | 0.25 | No |  | mosquitoes per visit | 1 | 4 | 0.25 | Yes |
| VIlag2 | 1 | 3 | 0.33 | No |  | VIlag4 | 4 | 3 | 1.33 | No |  | bldg footprint peri total | 1 | 4 | 0.25 | No |  | mosquitoes per visit | 1 | 4 | 0.25 | Yes |
| VIlag2 | 1 | 3 | 0.33 | No |  | adult obs per visit | 1 | 4 | 0.25 | Yes |  | bldg footprint peri total | 1 | 4 | 0.25 | No |  | mosquitoes per visit | 1 | 4 | 0.25 | Yes |
| VIlag2 | 1 | 3 | 0.33 | No |  | adult obs per visit | 1 | 4 | 0.25 | Yes |  | bldg footprint peri total | 1 | 4 | 0.25 | No |  | mosquitoes per visit | 1 | 4 | 0.25 | Yes |
| VIlag2 | 1 | 3 | 0.33 | No |  | adult obs per visit | 1 | 4 | 0.25 | Yes |  | child obs per visit | 1 | 4 | 0.25 | Yes |  | mosquitoes per visit | 1 | 4 | 0.25 | Yes |
| VIlag2 | 1 | 3 | 0.33 | No |  | adult obs per visit | 1 | 4 | 0.25 | Yes |  | child obs per visit | 1 | 4 | 0.25 | Yes |  | mosquitoes per visit | 1 | 4 | 0.25 | Yes |
| VIlag2 | 1 | 3 | 0.33 | No |  | adult obs per visit | 1 | 4 | 0.25 | Yes |  | child obs per visit | 1 | 4 | 0.25 | Yes |  | mosquitoes per visit | 1 | 4 | 0.25 | Yes |
| VIlag2 | 1 | 3 | 0.33 | No |  | adult obs per visit | 1 | 4 | 0.25 | Yes |  | child obs per visit | 1 | 4 | 0.25 | Yes |  | mosquitoes per visit | 1 | 4 | 0.25 | Yes |
| VIlag2 | 1 | 3 | 0.33 | No |  | adult obs per visit | 1 | 4 | 0.25 | Yes |  | child obs per visit | 1 | 4 | 0.25 | Yes |  | NDVI | 1 | 4 | 0.25 | No |
| VIlag2 | 1 | 3 | 0.33 | No |  | adult obs per visit | 1 | 4 | 0.25 | Yes |  | child obs per visit | 1 | 4 | 0.25 | Yes |  | NDVI | 1 | 4 | 0.25 | No |
| Vilag2 | 1 | 3 | 0.33 | No |  | adult obs per visit | 1 | 4 | 0.25 | Yes |  | child obs per visit | 1 | 4 | 0.25 | Yes |  | NDVI | 1 | 4 | 0.25 | No |
| VIlag3 | 2 | 3 | 0.67 | No |  | avg bldg area: avg lot area | 1 | 4 | 0.25 | No |  | child obs per visit | 1 | 4 | 0.25 | Yes |  | NDVI | 1 | 4 | 0.25 | No |
| VIlag3 | 2 | 3 | 0.67 | No |  | avg bldg area: avg lot area | 4 | 4 | 1.00 | No |  | child obs per visit | 1 | 4 | 0.25 | Yes |  | NDVI | 1 | 4 | 0.25 | No |
| VIlag3 | 2 | 3 | 0.67 | No |  | avg bldg area: avg lot area | 3 | 4 | 0.75 | No |  | Culex per visit | 1 | 4 | 0.25 | Yes |  | senior obs per visit | 1 | 4 | 0.25 | Yes |
| VIlag3 | 2 | 3 | 0.67 | No |  | avg bldg area: avg lot area | 3 | 4 | 0.75 | No |  | Culex per visit | 1 | 4 | 0.25 | Yes |  | senior obs per visit | 1 | 4 | 0.25 | Yes |
| VIlag3 | 2 | 3 | 0.67 | No |  | avg bldg area: avg lot area | 4 | 4 | 1.00 | No |  | Culex per visit | 1 | 4 | 0.25 | Yes |  | senior obs per visit | 1 | 4 | 0.25 | Yes |
| VIlag3 | 2 | 3 | 0.67 | No |  | avg bldg area: avg lot area | 1 | 4 | 0.25 | No |  | Culex per visit | 1 | 4 | 0.25 | Yes |  | senior obs per visit | 1 | 4 | 0.25 | Yes |
| VIlag3 | 2 | 3 | 0.67 | No |  | avg bldg peri: avg lot area | 1 | 4 | 0.25 | No |  | Culex per visit | 1 | 4 | 0.25 | Yes |  | senior obs per visit | 1 | 4 | 0.25 | Yes |
| VIlag3 | 1 | 3 | 0.33 | No |  | bldg footprint area avg | 4 | 4 | 1.00 | No |  | Culex per visit | 1 | 4 | 0.25 | Yes |  | senior obs per visit | 1 | 4 | 0.25 | Yes |
| VIlag3 | 1 | 3 | 0.33 | No |  | bldg footprint area avg | 4 | 4 | 1.00 | No |  | Culex per visit | 1 | 4 | 0.25 | Yes |  | senior obs per visit | 1 | 4 | 0.25 | Yes |
| VIlag3 | 1 | 3 | 0.33 | No |  | bldg footprint area avg | 4 | 4 | 1.00 | No |  | Culex per visit | 1 | 4 | 0.25 | Yes |  | senior obs per visit | 1 | 4 | 0.25 | Yes |
| VIlag3 | 1 | 3 | 0.33 | No |  | bldg footprint area avg | 4 | 4 | 1.00 | No |  | Culex per visit | 1 | 4 | 0.25 | Yes |  | total bldg area: total lot area | 1 | 4 | 0.25 | No |
| VIlag3 | 1 | 3 | 0.33 | No |  | bldg footprint area avg | 4 | 4 | 1.00 | No |  | Culex per visit | 1 | 4 | 0.25 | Yes |  | total bldg area: total lot area | 1 | 4 | 0.25 | No |
| VIlag3 | 2 | 3 | 0.67 | No |  | bldg footprint area avg | 4 | 4 | 1.00 | No |  | Culex per visit | 1 | 4 | 0.25 | Yes |  | total bldg area: total lot area | 1 | 4 | 0.25 | No |
| VIlag4 | 1 | 3 | 0.33 | No |  | bldg footprint area avg | 4 | 4 | 1.00 | No |  | female obs per visit | 1 | 4 | 0.25 | Yes |  | total bldg area: total lot area | 1 | 4 | 0.25 | No |
| VIlag4 | 4 | 3 | 1.33 | No |  | bldg footprint area avg | 4 | 4 | 1.00 | No |  | female obs per visit | 1 | 4 | 0.25 | Yes |  | total bldg area: total lot area | 1 | 4 | 0.25 | No |
|  |  |  |  |  |  |  |  |  |  |  |  |  |  |  |  |  |  | total bldg peri: total lot area | 1 | 4 | 0.25 | No |
|  |  |  |  |  |  |  |  |  |  |  |  |  |  |  |  |  |  | total obs per visit | 1 | 4 | 0.25 | Yes |
