## Supplemental Table 4 for "Dynamics of data availability in disease modeling: An example evaluating the trade-offs of ultra-fine-scale factors applied to human West Nile virus disease models in the Chicago area, USA"

| **Regression Method^a^** | **Model** | **Included Covariates** | **N covariates** | ***P*** | **R^2^** | **ROC^b^** | **BIC** | **Δ BIC** |
| --- | --- | --- | --- | --- | --- | --- | --- | --- |
| Logistic | MIR & Mosquito Abundance  (E1C201) | - tempc - preci + templag1 + templag3^*^ - precilag1^*^ - precilag3 - precilag4 + mir mean - mir diff + mirlag1 + mirlag2 + mirlag3^**^ + mirlag4^*^ + totpop + blackpct + asianpct + dmipct - dhipct + ccpct - hpctpreww - hpctpostww - hpct7089 + abund - abundlag1 + abundlag2 + abundlag3 + abundlag4 + bldg footprint area avg - bldg footprint peri avg + resi lot peri total - resi lot peri avg - avg bldg area:avg lot area - total bldg peri:total lot area | 34 | <0.0001 | 0.1652 | 0.88839 | 768.7 | 128.3 |
|  | Vector Index  (E2C2O1) | - tempc - preci + templag1 + templag2 + templag3^**^ - precilag1^*^ - precilag4 + totpop + blackpct + asianpct + hispanicpct - Income + dospct - dmipct - hpctpreww - hpct7089 + CB + avg bldg area:avg lot area^***^ + VI - VIlag1 + VIlag2 + VIlag3^*^ + VIlag4 | 23 | <0.0001 | 0.1382 | 0.8606 | 661.1 | 20.7 |
|  | Optimal  (E3C2O1) | - tempc - preci + templag1 + templag2 + templag3^**^ - precilag1^*^ - precilag3 + mirlag3^**^ + totpop - whitepct + blackpct - Income + dospct + dmipct - hpctpreww - hpct7089 - abundlag1 - bldg footprint area total + avg bldg area:avg lot area^**^ + VIlag4^*^ | 21 | <0.0001 | 0.1424 | 0.85941 | 640.4 | 0 |
|  | Global  (excluded) | - tempc^*^ - preci^*^ - yr + templag1^*^ + templag2 + templag3 - templag4 - precilag1^**^ - precilag2 - precilag3 - precilag4^*^ + MIRmean - MIRdiff + mirlag1 + mirlag2 + mirlag3^*^ + mirlag4 + totpop - whitepct - blackpct - asianpct - hispanicpct + income - owpct - dospct - dlipct - dmipct - dhipct - blpct - dfpct - mfpct - shrubpct - glandpct - pasturepct - wwpct - dtotpct - ftotpct + wtotpct - Jantemp + hpctpreww + hpctpostww + hpct7089 + hpctpost90 - abund - abundlag1 - abundlag2 + abundlag3 + abundlag4 + CB - bldg footprint total area + bldg footprint area avg - bldg footprint peri total - bldg footprint peri avg + resi lot area total + resi lot area avg + resi lot peri total - resi lot peri avg + total bldg area:total lot area + VI - VIlag1 - VIlag2 - VIlag3 - VIlag4 + Light pol + NDVI | 65 | 0.0009 | 0.3679 | 0.97097 | 833.492 | 193.092 |
| Linear | MIR & Mosquito Abundance  (E1C2O2) | mir mean + mir diff + mirlag1 + mirlag2 + mirlag3 + mirlag4^*^ + blpct^**^ + abund - abundlag1 - abundlag2 + abundlag3 + abundlag4^*^ + bldg footprint area avg^***^ | 13 | <0.0001 | 0.00578 | N/A | -182037 | 3358 |
|  | Vector Index  (E2C2O2) | + bldg footprint area avg^***^ + avg bldg area:avg lot area^***^ + VI - VIlag1 + VIlag2 + VIlag3^*^ + VIlag4^***^ | 7 | <0.0001 | 0.007271 |  | -185373 | 22 |
|  | Optimal  (E3C2O2) | - mirlag4 + bldg footprint area avg^** *^+ avg bldg area:avg lot area^**^ + VIlag4^***^ | 4 | <0.0001 | 0.006756 |  | -185395 | 0 |
|  | Global  (excluded) | - tempc - preci + yr - templag1 + templag2 + templag3 + templag4 - precilag1 + precilag2 - precilag3 - precilag4 + MIRmean - MIRdiff + mirlag1 - mirlag2 + mirlag3 - mirlag4^**^ - totpop - whitepct + blackpct + asianpct + hispanicpct + income + owpct - dospct - dlipct - dmipct - dhipct - blpct - dfpct - mfpct + shrubpct - glandpct + pasturepct + wwpct - dtotpct - ftotpct + wtotpct + Jantemp - hpctpreww - hpctpostww - hpct7089 - hpctpost90 - abund - abundlag1 - abundlag2 + abundlag3 + abundlag4 + CB - bldg footprint total area - bldg footprint area avg + bldg footprint peri total + bldg footprint peri avg - resi lot area total - resi lot area avg + resi lot peri total - resi lot peri avg + total bldg area:total lot area + - avg bldg area:avg lot area - # blgds - VI - VIlag1 - VIlag2 - VIlag3 + VIlag4^***^ + Light pol + NDVI | 68 | 0.0044 | 0.017482 |  | -94558.2 | 90836.8 |

A.

Table S4. Model fit comparisons of the UFS hexagons, applying (A) newly acquired data (excluding HLC and human observations, covariate set 2), or (B) only the covariates made available to the previously published Cook & DuPage model (covariate set 4). Each model outcome was assessed using logistic (presence/absence WNV human illness case) and generalized linear (WNV case rates, controlling for human population) methods. Asterisks indicate level of statistical significance (* = p ≤ 0.05, ** = p ≤ 0.001, *** = p ≤ 0.0001

^a^Logistic regression outcome = human WNV presence/absence per hexagon, per week; GLM outcome = WNV human case rate (per hexagon, per week).

^b^ROC applies to only logistic regression

^c^As the final selected model in the Original Cook & DuPage paper (2019), this model environment was assessed only for the comparison to the Cook & DuPage models for this study and not applied to the UFS model. The original model covariates, eftpct and ehwpct, have 0 observations among the selected 55 hexagons and were removed.

| **Regression Method^a^** | **Model** | **Included Covariates** | **N covariates** | ***P*** | **R^2^** | **ROC^b^** | **BIC** | **Δ BIC** |
| --- | --- | --- | --- | --- | --- | --- | --- | --- |
| Logistic | Optimal Cook & DuPage^+^  (E0C4O1) | - Yr - templag2 + templag3^*^ + templag4^*^ - Jantemp + mirlag1 + mirlag2 + mirlag3 + mirlag4^*^ + totpop - owpct - dlipct - dfpct - glandpct + hpctpost90 | 15 | <0.0001 | 0.1227 | 0.84897 | 632.3 | 56.1 |
|  | MIR & Mosquito Abundance  (E1C4O1) | dhipct - mfpct - wwpct + mirlag1 + mirlag2 + mirlag3^*^ + mirlag4 + templag3 + templag4^*^ - precilag1^*^ - precilag2 - precilag4^*^ + asianpct^**^ + totpop + mir mean - mir diff +abund - abundlag1^*^ + abundlag2 + abundlag3 + abundlag4 | 21 | <0.0001 | 0.1466 | 0.87537 | 653.3 | 77.1 |
|  | Vector Index  (E2C4O1) | dhipct - mfpct - wwpct + templag3^*^ + templag4^*^ - precilag1^*^ - precilag2 - precilag4^*^ + asianpct + VIlag1 + VIlag2 + VIlag3^*^ + VIlag4 | 14 | <0.0001 | 0.1404 | 0.85801 | 576.2 | 0 |
|  | Optimal 55 hex fitted  (E3C4O1) | dhipct - mfpct - wwpct + mirlag3^*^ + mirlag4 + templag3^*^ + templag4^**^ - precilag1^*^ - precilag2 - precilag4^*^ + asianpct^**^ + totpop | 12 | <0.0001 | 0.1646 | 0.89076 | 580.8 | 4.6 |
|  | Global  (excluded) | Yr - dospct - dlipct - dmipct - dmipct - dhipct - dfpct - mfpct + blpct - shrubpct - glandpct - pasturepct - ccpct - wwpct - owpct + mirlag1 + mirlag2 + mirlag3^*^ + mirlag4 - templag1 - templag2 + templag3^*^ + templag4^*^ - precilag1^*^ - precilag2 - precilag3 - precilag4^*^ + whitepct - blackpct - asianpct + hispanicpct - Income - totpop + Jantemp | 33 | <0.0001 | 0.179 | 0.89502 | 792 | 215.8 |
| Linear | Optimal Cook & DuPage  (E0C4O2) | Yr - templag2 + templag3 + templag4 + Jantemp + mirlag1 + mirlag2 + mirlag3^*^ + mirlag4^**^ - totpop^*^ - owpct^*^ - dlipct - dfpct^*^ - glandpct + hpctpost90 | 15 | <0.0001 | 0.004596 | N/A | -227354 | 90 |
|  | MIR & Mosquito Abundance  (E1C4O2) | dmipct^**^ + blpct^**^ + mirlag1 + mirlag2 + mirlag3 + mirlag4^*^ + templag3 - precilag1 - precilag4 - totpop^*^ - mir mean + mir diff - abund - abundlag1 + abundlag2 - abundlag3 + abundlag4^*^ | 17 | <0.0001 | 0.005914 |  | -182001 | 45443 |
|  | Vector Index  (E2C4O2) | dlipct + dmipct^***^ + templag3^*^ - totpop^**^ - VI - VIlag1 + VIlag2 + VIlag3^*^ + VIlag4^***^ | 9 | <0.0001 | 0.006639 |  | -185347 | 42097 |
|  | Optimal 55 hex fitted  (E3C4O2) | dmipct^**^ + blpct^**^ + mirlag3^**^ + mirlag4^**^ + templag3^*^ - precilag1 - precilag4 - totpop^*^ | 8 | <0.0001 | 0.004999 |  | -227444 | 0 |
|  | Global  (excluded) | Yr - dospct - dlipct - dmipct - dmipct - dhipct - dfpct - mfpct + blpct - shrubpct - glandpct - pasturepct - ccpct - wwpct - owpct + mirlag1 + mirlag2 + mirlag3^*^ + mirlag4^*^ - templag1 - templag2 + templag3 + templag4 - precilag1 - precilag2 - precilag3 - precilag4 + whitepct - blackpct - asianpct + hispanicpct - Income - totpop^*^ + Jantemp | 33 | <0.0001 | 0.005739 |  | -227199 | 245 |

Table S4 (continued).

B.
