## Supplemental Table 5 for "Dynamics of data availability in disease modeling: An example evaluating the trade-offs of ultra-fine-scale factors applied to human West Nile virus disease models in the Chicago area, USA"

| **Regression Type** | **Model** | **Included Covariates** | **N covariates** | ***P*** | **R^2^** | **ROC** | **BIC** | **Δ BIC** |
| --- | --- | --- | --- | --- | --- | --- | --- | --- |
| Logistic | MIR & Mosquito Abundance  (E1C1O1) | - tempc - preci + templag1 - templag2 + templag3* - precilag1* - precilag3 - precilag4 + mirmean - mirdiff + mirlag1 + mirlag2 + mirlag3** + mirlag4 - whitepct + blackpct + dospct - dmipct - hpctpreww - hpct7089 + abund - abundlag1 + abundlag2 + abundlag3 + abundlag4 + bldg footprint area total - total bldg area:total lot area + male obs per visit + female obs per visit + *Culex* per visit | 30 | <0.0001 | 0.1443 | 0.86736 | 742.5 | 107.9 |
|  | Vector Index  (E2C1O1) | - tempc - preci + templag1 + templag2 + templag3* - precilag1* - precilag4 + mirlag1 + mirlag3 - whitepct + blackpct - dospct + dmipct - hpctpreww - hpctpostww - hpct7089 - abund + Light pol + male obs per visit + female obs per visit + *Culex* per visit + VI - VIlag1 + VIlag2 + VIlag3 + VIlag4 | 26 | <0.0001 | 0.1311 | 0.85958 | 692.7 | 58.1 |
|  | Optimal  (E3C1O1) | - tempc - preci + templag1 + templag2 + templag3^**^ - precilag1^*^ - precilag4 + mirlag3^**^ + blackpct - dospct + dmipct - hpctpreww - hpct7089 + abund + Light pol + male obs per visit + female obs per visit + *Culex* per visit + VIlag4 | 19 | <0.0001 | 0.1171 | 0.838 | 634.6 | 0 |
|  | Global^a^  (excluded) | - tempc* - preci* - yr + templag1* + templag2 + templag3 - templag4 - precilag1** - precilag2 - precilag3 - precilag4* + mirmean - mirdiff + mirlag1 + mirlag2 + mirlag3* + mirlag4 - totpop + whitepct + blackpct + asianpct + hispanicpct + income - owpct - dospct - dlipct - dmipct - dhipct - blpct - dfpct - mfpct - shrubpct - glandpct - pasturepct + wwpct - dtotpct - ftotpct + wtotpct - Jantemp + hpctpreww + hpctpostww + hpct7089 + hpctpost90 - abund - abundlag1 - abundlag2 - abundlag3 + abundlag4 + CB - bldg footprint total area - bldg footprint area avg + bldg footprint peri total + bldg footprint peri avg - resi lot area total - resi lot area avg - resi lot peri total + resi lot peri avg - total bldg area:total lot area - # blgds + VI - VIlag1 - VIlag2 - VIlag3 - VIlag4 + Light pol + NDVI + mosquitoes per visit | 65 | 0.0009 | 0.3679 | 0.97097 | 772.8 | 138.2 |
| Linear | MIR & Mosquito Abundance  (E1C1O2) | mir mean + mir diff + mirlag1 + mirlag2 + mirlag3 + mirlag4^*^ + blpct^**^ + abund - abundlag1 - abundlag2 + abundlag3 + abundlag4^*^ + bldg footprint area avg^***^ | 13 | <0.0001 | 0.00578 | N/A | -182037 | 3352 |
|  | Vector Index  (E2C1O2) | templag3* + bldg footprint area avg^***^ - resi lot peri avg + VI - VIlag1 + VIlag2 + VIlag3^*^ + VIlag4^***^ | 8 | <0.0001 | 0.007178 |  | -185362 | 27 |
|  | Optimal  (E3C1O2) | - mirlag4 + bldg footprint area avg^***^- resi lot peri avg + VIlag4^***^ | 4 | <0.0001 | 0.006236 |  | -185389 | 0 |
|  | Global^b^  (excluded) | - tempc - preci + yr - templag1 + templag2 + templag3 + templag4 - precilag1 + precilag2 - precilag3 - precilag4 + mirmean - mirdiff + mirlag1 - mirlag2 + mirlag3 - mirlag4^**^ - totpop + whitepct + blackpct + asianpct + hispanicpct + income - owpct - dospct - dlipct - dmipct - dhipct - blpct - dfpct - mfpct - shrubpct - glandpct - pasturepct + wwpct - dtotpct - ftotpct + wtotpct + Jantemp - hpctpreww - hpctpostww - hpct7089 - hpctpost90 - abund - abundlag1 - abundlag2 - abundlag3 + abundlag4 + CB - bldg footprint total area - bldg footprint area avg + bldg footprint peri total + bldg footprint peri avg - resi lot area total - resi lot area avg + resi lot peri total - resi lot peri avg - total bldg area:total lot area - avg bldg area:avg lot area - # blgds - VI - VIlag1 - VIlag2 - VIlag3 + VIlag4^***^ + Light pol + NDVI + senior obs per visit | 64 | 0.0022 | 0.017241 |  | -94591.4 | 90797.6 |

A.

| **Regression Type^a^**  B. | **Model** | **Included Covariates** | **N covariates** | ***P*** | **R^2^** | **ROC^b^** | **AICc** | **Δ AICc** |
| --- | --- | --- | --- | --- | --- | --- | --- | --- |
| Logistic | Optimal Cook & DuPage  (E0C3O1) | - Yr - templag2 + templag3^*^ + templag4^*^ - Jantemp + mirlag1 + mirlag2 + mirlag3* + mirlag4^*^ - totpop - owpct - dlipct - dfpct - glandpct + hpctpost90 - senior obs per visit - adult obs per visit + child obs per visit + male obs per visit - mosquitoes per visit + *Culex* per visit | 21 | <0.0001 | 0.1337 | 0.86184 | 683.4 | 10.7 |
|  | MIR & Mosquito Abundance^c^  (E1C3O1) | - tempc - preci + templag1 + templag2 - precilag1* - precilag4* - mirmean* - mirdiff + mirlag1 + mirlag2 + mirlag3** - mirlag4* - totpop + blackpct + dospct + hpctpostww + hpct7089 + hpctpost90* + abund - abundlag1 + abundlag2 + abundlag3 + abundlag4 + bldg footprint peri avg + resi lot area total + avg bldg peri:avg lot area** + senior obs per visit + adult obs per visit + child obs per visit - male obs per visit + mosquitoes per visit + *Culex* per visit | 32 | <0.0001 | 0.1508 | 0.86968 | 757.7 | 61.1 |
|  | Vector Index  (E2C3O1) | - tempc - preci + templag1 + templag2 + templag3** - precilag1* - precilag4 - whitepct + blackpct - dospct + dmipct + hpctpostww + hpct7089 + hpctpost90 + resi lot area total - senior obs per visit + adult obs per visit + child obs per visit - male obs per visit - mosquitoes per visit + *Culex* per visit + VI - VIlag1 + VIlag2 + VIlag3* + VIlag4 | 26 | <0.0001 | 0.1234 | 0.84561 | 696.6 | 23.9 |
|  | Optimal 55 hex fitted^d^  (E3C3O1) | - tempc - preci + templag1 + templag2 + templag3** - precilag1* - precilag4 - whitepct + blackpct - dospct + dmipct + hpctpostww + hpct7089 + resi lot area total - senior obs per visit + adult obs per visit + child obs per visit - male obs per visit - mosquitoes per visit + *Culex* per visit + VIlag4 + mirlag3** - abund | 23 | <0.0001 | 0.1153 | 0.834 | 672.7 | 0 |
|  | Global  (excluded) | Model Failed to Converge | N/A | | | | | |
| Linear | Optimal Cook & DuPage^e^  (E0C3O2) | Yr - templag2 + templag3 + templag4 + Jantemp + mirlag1 + mirlag2 + mirlag3^*^ + mirlag4^**^ - totpop^*^ - owpct^*^ - dlipct - dfpct^*^ - glandpct + hpctpost90 - senior obs per visit + adult obs per visit + child obs per visit - male obs per visit + mosquitoes per visit - *Culex* per visit | 21 | <0.0001 | 0.0048 | N/A | -227300 | 0 |
|  | MIR & Mosquito Abundance^e^  (E1C3O2) | blpct** + bldg footprint area avg*** - senior obs per visit + adult obs per visit + child obs per visit - male obs per visit - mosquitoes per visit + *Culex* per visit + mirmean + mirdiff + mirlag1 + mirlag2 + mirlag3 + mirlag4* + abund - abundlag1 - abundlag2 - abundlag3 + abundlag4* | 19 | <0.0001 | 0.005886 |  | -181982 | 45318 |
|  | Vector Index^f^  (E2C3O2) | bldg footprint area avg*** + adult obs per visit + child obs per visit + female obs per visit - total obs per visit - mosquitoes per visit + *Culex* per visit + VI - VIlag1 + VIlag2 + VIlag3* + VIlag4*** | 12 | <0.0001 | 0.006877 |  | -185322 | 41978 |
|  | Optimal 55 hex fitted^g^  (E3C3O2) | bldg footprint area avg*** - senior obs per visit - adult obs per visit + child obs per visit + female obs per visit - mosquitoes per visit + *Culex* per visit + VIlag4*** - mirlag4 | 9 | <0.0001 | 0.00635 |  | -185344 | 41956 |
|  | Global^g^  (excluded) | - preci + Yr - templag1 + templag2 + templag3 + templag4 - precilag1 + precilag2 - precilag3 - precilag4 - totpop + whitepct + blackpct + asianpct + hispanicpct - Income - owpct - dospct - dlipct - dmipct - dhipct + blpct - dfpct - mfpct - shrubpct + glandpct - pasturepct* - wwpct - dtotpct - ftotpct + wtotpct + Jantemp - hpctprewww - hpct7089 - hpctpost90 + CB - bldg footprint area total - bldg footprint area avg + bldg footprint peri total + lightpol + NDVI + senior obs per visit - adult obs per visit - child obs per visit + female obs per visit - mosquitoes per visit + *Culex* per visit - VI - VIlag1 - VIlag2 - VIlag3 + VIlag4*** + mirmean - mirdiff + mirlag1 - mirlag2 + mirlag3 - mirlag4** - abund - abundlag1 - abundlag2 - abundlag3 + abundlag4 | 63 | 0.0013 | 0.017472 |  | -94601.5 | 132698.5 |
