## Supplemental Table 6 for "Dynamics of data availability in disease modeling: An example evaluating the trade-offs of ultra-fine-scale factors applied to human West Nile virus disease models in the Chicago area, USA"

| **Predictor** | **Mean** | **Standard Error** |  | **Predictor** | **Mean** | **Standard Error** |
| --- | --- | --- | --- | --- | --- | --- |
| tempc | 20.42 | 0.0341 |  | hpctpost90 | 16.54 | 0.1312 |
| preci | 25.13 | 0.2354 |  | Trap_Mean | 15.76 | 0.1458 |
| templag1 | 20.12 | 0.0378 |  | Trap_Meanlag1 | 16.15 | 0.1507 |
| templag2 | 19.73 | 0.0418 |  | Trap_Meanlag2 | 16.47 | 0.1562 |
| templag3 | 19.17 | 0.0460 |  | Trap_Meanlag3 | 16.73 | 0.1627 |
| templag4 | 18.49 | 0.0502 |  | Trap_Meanlag4 | 16.92 | 0.1704 |
| precilag1 | 25.03 | 0.2310 |  | Trap_Sum | 8693.12 | 81.0305 |
| precilag2 | 25.13 | 0.2319 |  | CB | 153.31 | 0.9540 |
| precilag3 | 25.01 | 0.2316 |  | Water_Area | 366928.54 | 8649.3909 |
| precilag4 | 25.04 | 0.2281 |  | Senior_Obs per visit | 0.51 | 0.0057 |
| MIR_mean | 5.69 | 0.1090 |  | Adult_Obs per visit | 4.82 | 0.0209 |
| MIRLAG1_1 | 5.28 | 0.1071 |  | Child_Obs per visit | 2.02 | 0.0150 |
| MIRLAG2_1 | 4.82 | 0.1050 |  | Male_Obs per visit | 4.06 | 0.0188 |
| MIRLAG3_1 | 4.21 | 0.1020 |  | Female_Obs per visit | 3.30 | 0.0158 |
| MIRLAG4_1 | 3.41 | 0.0800 |  | Total Obs Visits | 6.76 | 0.0203 |
| totpop | 1371.58 | 7.4872 |  | Total_Obs per visit | 7.35 | 0.0334 |
| whitepct | 78.90 | 0.1232 |  | HLC Mosquitoes Total | 4.11 | 0.0406 |
| blackpct | 2.23 | 0.0252 |  | Mosquitos per visit | 0.55 | 0.0050 |
| asianpct | 10.14 | 0.0719 |  | Culex per visit | 0.12 | 0.0020 |
| hispanicpct | 16.36 | 0.1472 |  | Nuissance Factor observed | 5.33 | 0.0576 |
| Income | 78422.97 | 218.7765 |  | WNV Added Risk observed | 0.17 | 0.0025 |
| owpct | 2.36 | 0.0684 |  | bldg_footprint_area_total | 92796.82 | 326.2624 |
| dospct | 9.54 | 0.0719 |  | bldg_footprint_area_avg | 325.22 | 3.7038 |
| dlipct | 51.36 | 0.2031 |  | Building_Footprint_peri_total | 24686.05 | 102.2540 |
| dmipct | 22.06 | 0.1332 |  | Building_Footprint_peri_avg | 66.39 | 0.2376 |
| dhipct | 7.09 | 0.0785 |  | Residential_lot_area_total | 504061.25 | 1131.9698 |
| blpct | 0.02 | 0.0014 |  | Residential_lot_area_avg | 32852.08 | 1022.9782 |
| dfpct | 3.06 | 0.1072 |  | Residential_lot_peri_total | 47293.98 | 198.4891 |
| efpct | 0.00 | 0.0000 |  | Residential_lot_peri_avg | 2908.47 | 99.9071 |
| mfpct | 0.83 | 0.0243 |  | total_bldg_area/total_lot_area | 4.11 | 0.1572 |
| shrubpct | 0.46 | 0.0294 |  | avg_bldg_area/avg_lot_area | 0.15 | 0.0008 |
| glandpct | 0.99 | 0.0460 |  | total_bldg_peri/total_lot_area | 0.94 | 0.0378 |
| pasturepct | 0.83 | 0.0308 |  | avg_bldg_peri/avg_lot_area | 0.04 | 0.0002 |
| ccpct | 0.76 | 0.0332 |  | buildings | 444.24 | 2.4067 |
| wwpct | 0.78 | 0.0207 |  | bldg_density (sq.mi.) | 683.45 | 3.7027 |
| ehwpct | 0.00 | 0.0000 |  | persons_per_bldg | 3.82 | 0.0309 |
| dtotpct | 84.70 | 0.3311 |  | Vector Index | 8.49 | 0.2107 |
| ftotpct | 6.10 | 0.2078 |  | VIlag1 | 8.70 | 0.2199 |
| wtotpct | 1.06 | 0.0286 |  | VIlag2 | 8.73 | 0.2301 |
| Jantemp | -4.92 | 0.0269 |  | VIlag3 | 8.52 | 0.2391 |
| hpctpreww | 8.58 | 0.1034 |  | VIlag4 | 8.00 | 0.2385 |
| hpctpostww | 36.99 | 0.2073 |  | LightPollution | 5.04 | 0.0118 |
| hpct7089 | 37.90 | 0.2010 |  | NDVI | 0.50 | 0.0009 |
